# kallisto, bustools, and kb-python for quantifying bulk, single-cell, and single-nucleus RNA-seq

**DOI:** 10.1101/2023.11.21.568164

**Authors:** Delaney K. Sullivan, Kyung Hoi (Joseph) Min, Kristján Eldjárn Hjörleifsson, Laura Luebbert, Guillaume Holley, Lambda Moses, Johan Gustafsson, Nicolas L. Bray, Harold Pimentel, A. Sina Booeshaghi, Páll Melsted, Lior Pachter

## Abstract

The term “RNA-seq” refers to a collection of assays based on sequencing experiments that involve quantifying RNA species from bulk tissue, from single cells, or from single nuclei. The kallisto, bustools, and kb-python programs are free, open-source software tools for performing this analysis that together can produce gene expression quantification from raw sequencing reads. The quantifications can be individualized for multiple cells, multiple samples, or both. Additionally, these tools allow gene expression values to be classified as originating from nascent RNA species or mature RNA species, making this workflow amenable to both cell-based and nucleus-based assays. This protocol describes in detail how to use kallisto and bustools in conjunction with a wrapper, kb-python, to preprocess RNA-seq data.

## Introduction

### Overview

The preprocessing^1,2^ step of RNA-seq^3^ experiments involves mapping reads to a reference genome or transcriptome, followed by gene expression or transcript abundance quantification.^4^ Many open-source tools exist for bulk RNA-seq preprocessing^5–14^ as well as single-cell RNA-seq preprocessing.^1,15–21^ kallisto^8^ introduced the pseudoalignment paradigm for improving the accuracy of alignment and reducing runtimes and memory footprint of bulk RNA-seq preprocessing and, with the development of bustools^22^, has been adapted for both single-cell RNA-seq quantification^1^ and single-nucleus RNA-seq quantification.^23^ The bustools suite of tools operates on the read mapping results of kallisto and processes them to generate quantification results, which may involve unique molecular identifier (UMI)^24,25^ collapsing and barcode error correction for single-cell and single-nucleus assays. While multiple steps are necessary to process input consisting of FASTQ sequencing files, a reference genome FASTA, and a GTF annotation^26,27^, to an output of quantifications using kallisto and bustools, these steps are greatly facilitated by the wrapper tool, kb-python. kb-python can extract reference transcriptomes from reference genomes and run kallisto and bustools in workflows optimal for each assay type. The kb-python tool simplifies the running of kallisto and bustools to the extent that all of this can be done in two steps: “kb ref” for generating a kallisto index from an annotated reference genome and “kb count” for mapping and quantification. Thus, kallisto, bustools, and kb-python make RNA-seq preprocessing efficient, modular, flexible, and simple.^1^

#### Box 1: Software tools and their description

Software tools:

- **kallisto**: Performs pseudoalignment to a reference transcriptome and stores the mapping results in a BUS file.^8^
- **bustools**: Processes the results in the BUS file to correct barcodes, deduplicate UMIs, and generate quantification files (e.g. count matrices).^1,22^
- **kb-python**: A wrapper around kallisto and bustools that facilitates usage of those tools and facilitates the generation of a reference transcriptome. The kallisto and bustools binaries come packaged in kb-python.

Installation:

**pip install kb_python**

Note: To install kallisto and bustools from source code rather than using the precompiled binaries or to install specific versions of the software, see **Supplementary Note 1**.

### Index construction

For RNA-seq read mapping, kallisto builds an *index* from a set of sequences, referred to as targets, representing the set of sequences that RNA-seq reads can be mapped to. In a standard analysis, these targets are usually transcript sequences (i.e. each individual target corresponds to one transcript). However, more generally, users can define targets from any sets of sequences they wish to map their sequencing reads against. Since kallisto is a tool that leverages pseudoalignment, the mapping procedure relies on read assignment, such that each read is deemed to be compatible with a certain set of targets, rather than standard alignment. The kallisto index is based on the Bifrost^28^ implementation of the colored de Bruijn graph^29^, which enables memory-efficient and rapid read assignment.

kb-python enables the construction of kallisto indices through the **kb ref** command (**Fig. 1**). Different types of kallisto indices can be built by specifying the --workflow argument in kb ref, which selects the type of index to be constructed. The default is **--workflow=standard**, which creates an index suitable for bulk and single-cell RNA-seq quantification. It creates an index built from only the coding DNA sequences (the usage of coding DNA here follows that of Ensembl^30^, i.e., the sequences of the mature transcripts wherein introns are not included as they have been spliced out). The index created by **--workflow=nac** (nac: nascent and coding DNA) contains both the coding DNA *and* the nascent transcripts. The nascent transcript sequences consist of the full gene (both exons and introns). This nac index is suitable for single-nucleus RNA-seq as there exists a high abundance of non-mature transcripts captured in nucleus-based sequencing assays.^31^ Additionally, this nac index should be used for analyses that require jointly modeling nascent and mature RNA species.^32–37^ For both the standard and nac index types, a user supplies a genome FASTA and GTF annotation, which kb-python uses to extract the relevant sequences. Finally, if one wishes to index a custom set of targets or of k-mers (**Supplementary Note 2**), one can use **--workflow=custom** which builds an index from a FASTA file containing the target sequences of interest to be supplied.

**Fig. 1:**
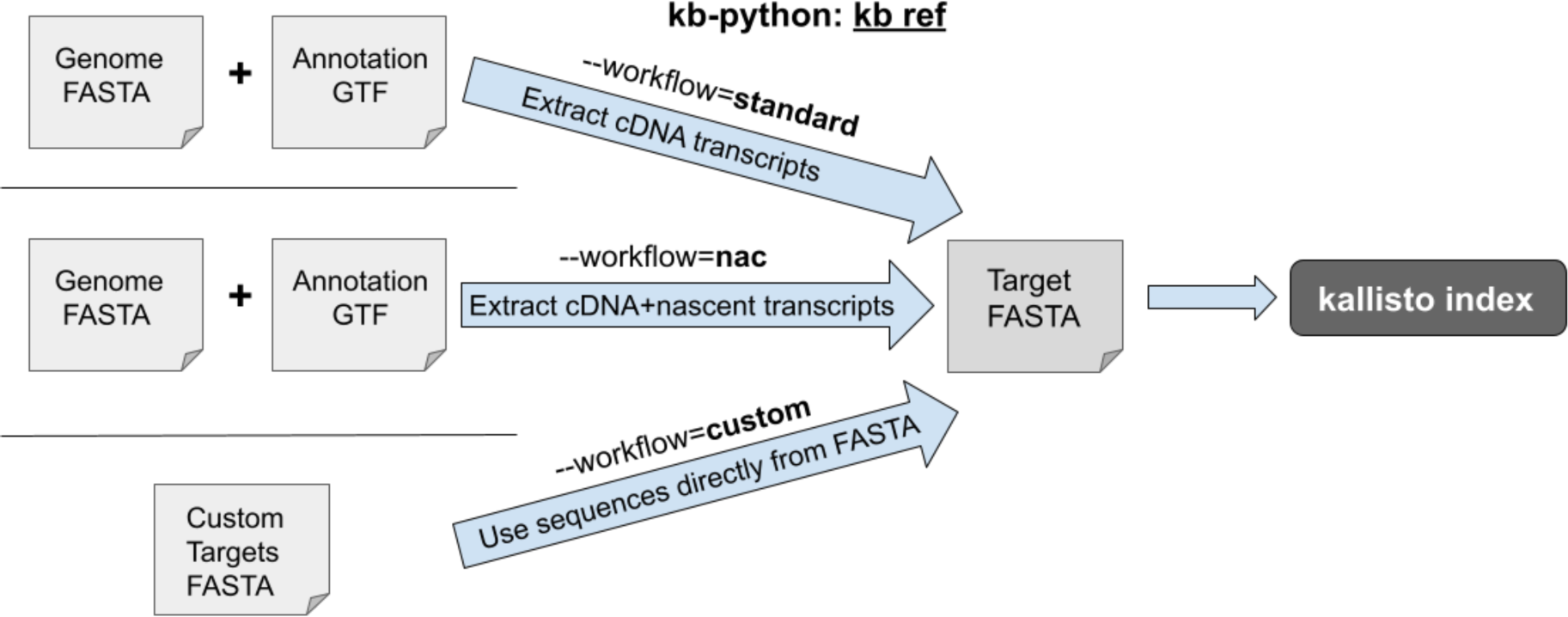
“kb ref” can be used to generate three different types of kallisto indices.

Creating the index in kb-python invokes the **kallisto index** command in the kallisto program (**Box 2**). Indexing with kb-python has the advantage that a reference transcriptome is generated directly from a FASTA and GTF ensuring consistency between the transcriptome reference, its associated index, and the input FASTA and GTF.

#### Box 2: kb ref

Below, we show how to run kb ref using three different index types. Only the underlined files need to be supplied by the user; the other files are output files generated as part of the indexing process and may be necessary for the subsequent mapping and quantification step. The corresponding kallisto index commands that are invoked are shown beneath each kb ref call (note that, by default, the kallisto index command is invoked using 8 threads).

1. **standard** index type (default): **kb ref -i index.idx -g t2g.txt -f1 cdna.fasta <u>genome.fasta genome.gtf</u>**

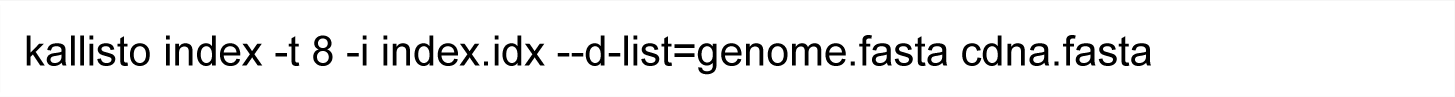

2. **nac** index type: **kb ref --workflow=nac -i index.idx -g t2g.txt -c1 cdna.txt -c2 nascent.txt \ -f1 cdna.fasta -f2 nascent.fasta <u>genome.fasta genome.gtf</u>**

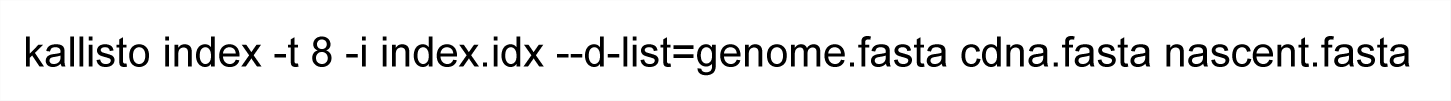

3. **custom** index type: **kb ref --workflow=custom -i index.idx <u>custom.fasta</u>**

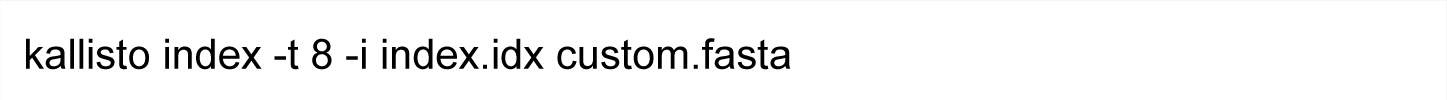

Explanation of output files:

- index.idx: The kallisto index that is generated
- t2g.txt: The transcript-to-gene mapping file
- cdna.fasta: The generated FASTA file containing the extracted coding DNA sequences
- nascent.fasta: The generated FASTA file with extracted nascent transcript sequences
- cdna.txt: The transcript names of the coding DNA sequences
- nascent.txt: The nascent transcript names (which are simply the gene names)

Additionally, using kb-python (via the --include-attributes and --exclude-attributes options) allows specific biotypes to be selected from the GTF file, making possible filtering of entries such as pseudogenes, which can improve read mapping accuracy^38^ and reduce memory usage (**Supplementary Note 3**). It is recommended to perform GTF filtering, especially for the nac index type where there will be many overlapping segments among annotated regions in the genome. While there is no universally defined best practice for GTF filtering, it is recommended that a user uses the CellRanger^39^ gene biotypes (shown in **Supplementary Note 3**) for standard single-cell and single-nucleus RNA-seq assays. More generally, if a user is unsure of what biotypes to include, it is recommended that the user selects only the specific biotypes that the user is interested in (e.g. selecting only protein coding genes if a user is only interested in protein coding genes). Finally, the kallisto index command has a --d-list option which improves the mapping specificity by isolating certain sequences, known as distinguishing flanking k-mers (DFKs), that may cause erroneous read mapping.^23^ The DFKs that are identified depend on the FASTA file supplied to the --d-list option. While the --d-list option can be entered by the user directly into kb ref, kb ref already by default calls kallisto index with the --d-list option set to the genome FASTA supplied but can be disabled by specifying --d-list=None in kb ref. For all analyses that involve RNA transcript quantification, it is recommended that the --d-list be set to the respective genome FASTA file to ensure good mapping specificity. This feature should typically only be used in any standard RNA-seq analysis (e.g. any usage with the standard index type or the nac index type produced by kb ref). This feature should not be used in other cases where custom non-transcript targets are indexed.

### Mapping and quantification

The **kb count** command within kb-python enables mapping and quantification of bulk, single-cell, and single-nucleus RNA-seq reads (**Fig. 2**). As different sequencing assays have different read structures, strandedness, parity, and barcodes, one must provide the specifications for the technology which produced the sequencing reads.

**Fig. 2:**
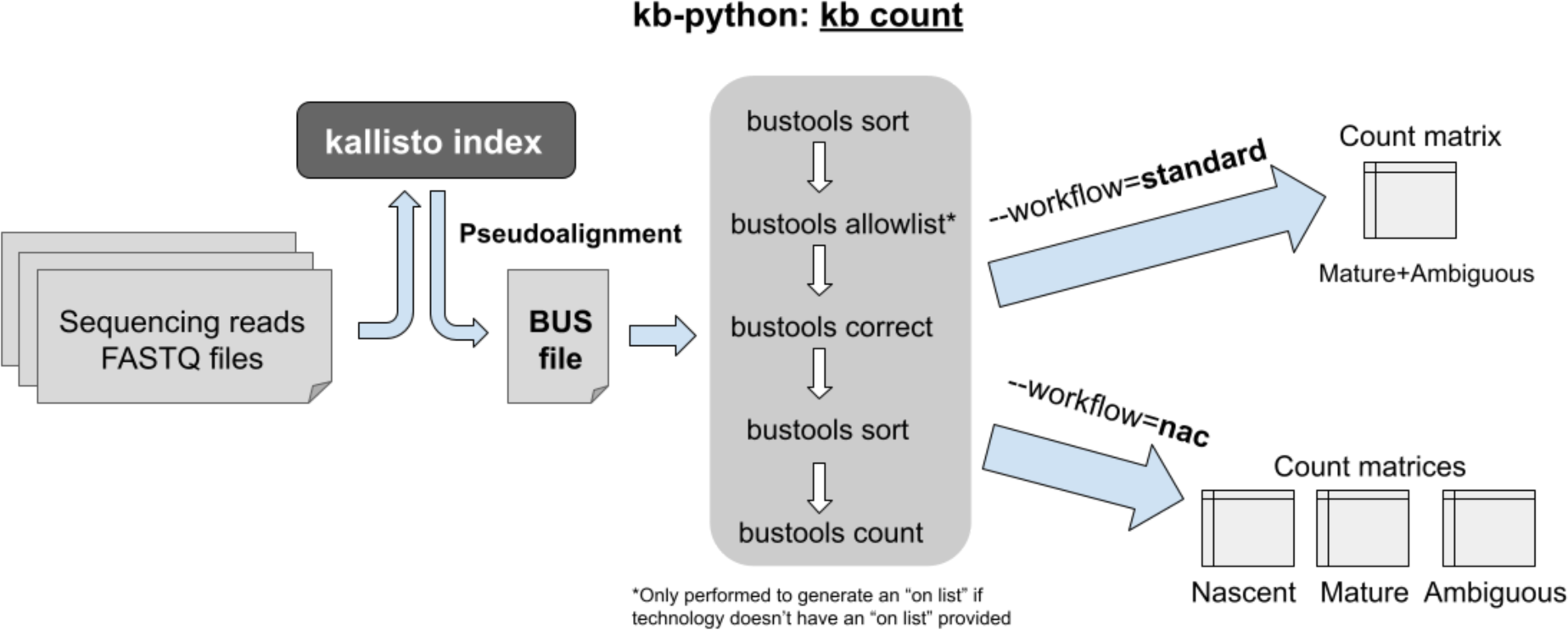
“kb count” can be used to produce quantifications in the form of count matrices for bulk, single-cell, and single-nucleus RNA-seq.

The specifications for sequencing assay technology within kb-python are as follows:

- **Technology string**: A *technology string* for a particular type of assay can be supplied via the **-x** option. The technology string can be used in one of three ways:

○ Option 1: Several assays are predefined within the software (the list is viewable by calling **kb --list**) so one can name one of those directly (e.g. one can specify -x 10xv3).
○ Option 2: One can use seqspec^40,41^ which contains machine-readable specifications for a wide range of sequencing assays.
○ Option 3: One can format their own custom technology string specifying the read locations of the barcodes, unique molecular identifiers (UMIs), and the biological sequence that is to be mapped (**Box 3**).
- **Strandedness**: If a read (or the first read in the case of paired-end reads) is to be mapped in forward orientation, one should specify **--strand=forward**. If it is to be mapped in reverse orientation, one should specify **--strand=reverse**. If one does not want to map reads with strand-specificity, then one should specify **--strand=unstranded**. If a predefined name is used in the technology string -x option (option 1), then kb-python uses a default stranded option for that technology (e.g. for 10xv3, the default is forward); otherwise, the default is unstranded. Setting the --strand option explicitly will overrule the default option.
- **Parity**: If the technology involves two biological read files that are derived from paired-end sequencing (as is the case with Smartseq2^42,43^ and Smartseq3^44^ and many bulk RNA sequencing kits), one should specify **--parity=paired** to perform mapping that takes into account the fact that the reads are paired-end. Otherwise, one can specify **--parity=single**. If a predefined name is used in the -x technology string option (option 1), then kb-python uses the default parity option for that technology (e.g for -x Smartseq2, --parity=paired is already enabled by default).
- **On list**: For single-cell and single-nucleus sequencing assays, barcodes are used to identify each cell or nucleus. The “on list” of barcodes represents the known barcode sequences that are included in the assay. Barcodes extracted from the sequencing reads will be error-tolerantly mapped to this list in a process known as barcode error correction. The on list filename can be specified with the **-w** option in kb count. It can also be obtained by seqspec.^40^ If an on list is not provided or cannot be found for the given technology, then an on list is created by bustools via the **bustools allowlist** command which identifies repeating barcodes in sequencing reads. If the technology does not include cell barcodes (as is the case in bulk RNA-seq), the “on list” option is irrelevant and no barcode processing occurs which should be the case for assays that don’t include cell/nuclei barcodes (skipping barcode error correction can also be done by specifying **-w NONE**). If a predefined name is used in the -x technology string option (option 1), then kb-python uses the default on list option for that technology.

#### Box 3: Custom technology string

The custom technology string (supplied to -x) contains the format **barcode:UMI:DNA**, representing the locational information of the barcode, UMI, and the DNA (where DNA is the biological read to be mapped):

**-x a,b,c:d,e,f:g,h,i**

- a: barcode file number, b: barcode start position, c: barcode end position
- d: UMI file number, e: UMI start position, f: UMI end position
- g: DNA file number, h: DNA start position, i: DNA end position

Important notes: File numbers and positions are zero-indexed. If no specific end position exists (i.e. the end position is the very end of the read), the end position should be set to 0. If cell barcodes and/or UMIs are not supported by the technology, the barcode and/or UMI field can be set to -1,0,0.

Thus, for 10xv3:

**-x 0,0,16:0,16,28:1,0,0**

Sequences can be stitched together by specifying multiple locations; for example, a SPLiT-seq^45^ assay, which contains three separate unlinked barcodes, each of length 8, and a UMI of length 10 in the second file and the DNA in the first file would look as follows:

**-x 1,10,18,1,48,56,1,78,86:1,0,10:0,0,0**

Final note about multiple locations: If the paired-end read mapping option is enabled, exactly two DNA locations should be specified (for the first and second read in the pair).

If a technology does not fit into this format (e.g. due to barcodes or UMIs of variable lengths and positions), preprocessing of the FASTQ file should be performed beforehand to reformat the reads into a structure that can be handled by this format.^46^

If a nac index was generated by kb ref, **--workflow=nac** should be used in kb count so that the nascent and mature RNA species are quantified accurately; otherwise that option should be omitted or **--workflow=standard** (which is the default) can be explicitly specified. For the nac index type, one obtains three count matrices: 1) nascent, 2) mature, and 3) ambiguous. In most experiments, the plurality of reads will be “ambiguous” since they originate from exons, which are present in both nascent RNA and mature RNA. Therefore, it is desirable to generate additional matrices by adding the counts from those three matrices, which users can either do themselves or by using the --sum option.^23^ **--sum=total** adds all three matrices, **--sum=cell** adds the mature and ambiguous matrices, and **--sum=nucleus** adds the nascent and ambiguous matrices. Different matrices may be used for different types of analyses. For example, in single-cell RNA-seq analysis (where most “ambiguous” counts are likely of mature RNA origin), jointly modeling the mature+ambiguous count matrix (--sum=cell) with the nascent count matrix permits biophysical modeling of RNA processing.^34,37^ In single-nucleus RNA-seq quantification, one might want to use --sum=nucleus to add up the nascent+ambiguous counts. The kb-python, kallisto, and bustools commands for the standard and nac index types are shown in **Box 4** and **Box 5**, respectively.

#### Box 4: kb count (standard index type)

Below, we show how to run kb count using the standard index type (which is the default used if no --workflow option is explicitly specified). The underlined files need to be supplied by the user; these include the files generated from the kb ref command as well as the FASTQ sequencing reads. The corresponding kallisto and bustools commands (as well as Unix commands to create and remove files/directories) that are called by kb count are shown beneath each kb count command (note that, by default, 8 threads and 2 gigabytes of memory are assigned).

**kb count -x <tech> -w <u>onlist.txt</u> -o output_dir -i <u>index.idx</u> -g <u>t2g.txt R1.fastq R2.fastq</u>**

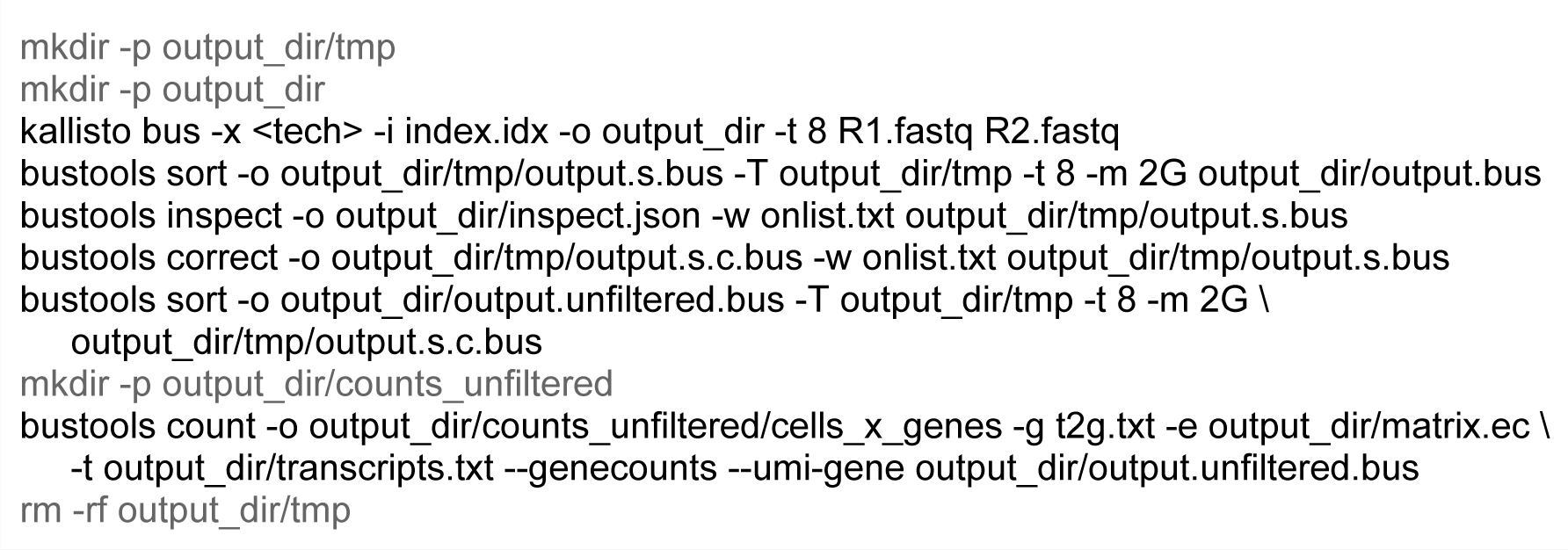

- <tech>: The technology string
- onlist.txt: The name of the file containing the “on list” of barcodes

○ Specify NONE to skip barcode error correction, or omit completely to have bustools create its own “on list” for correction

Note: In the workflow above, the following options in kb count can be used:

- **--parity=single or --parity=paired**
- **--strand=forward** or **--strand=reverse** or **--strand=unstranded**

One can alternatively set those options at the end of <tech>, e.g.: <tech>%forward%paired

The R1.fastq and R2.fastq inputs can be replaced with multiple sets of read files listed consecutively, as long as each pair is in order.

#### Box 5: kb count (nac index type)

Below, we show how to run kb count using the nac index type. The underlined files need to be supplied by the user;these include the files generated from the kb ref command using --workflow=nac as well as the FASTQ sequencing reads. The corresponding kallisto and bustools commands (as well as Unix commands to create and remove files/directories) that are called are shown beneath each kb count call (note that, by default, 8 threads and 4 gigabytes of memory are used).

**kb count -x <tech> --workflow=nac -w <u>onlist.txt</u> -o output_dir -i <u>index.idx</u> -g <u>t2g.txt</u> \ -c1 <u>cdna.txt</u> -c2 <u>nascent.txt</u> --sum= <sum> <u>R1.fastq R2.fastq</u>**

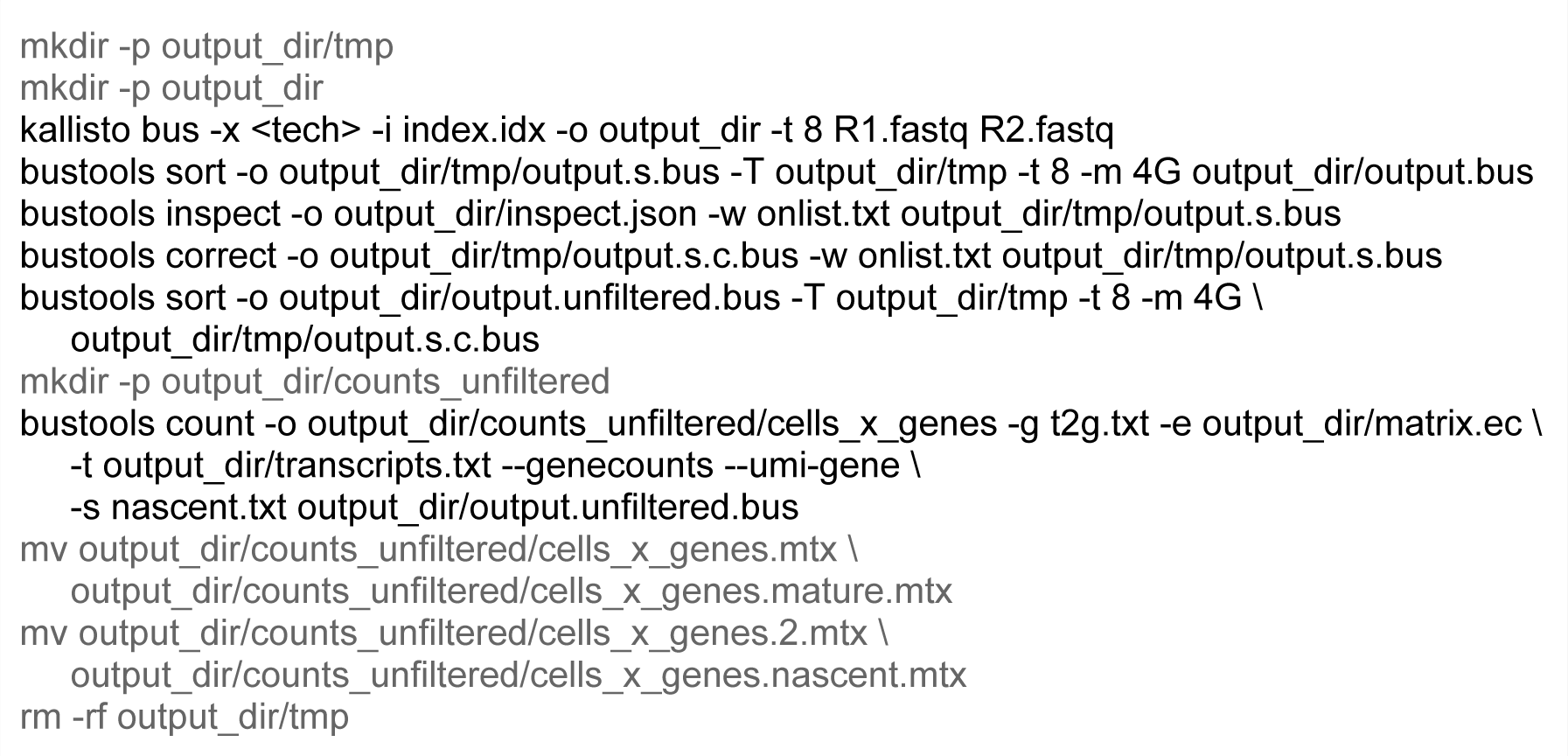

- <tech>: The technology string
- onlist.txt: The name of the file containing the “on list” of barcodes

○ Specify NONE to skip barcode error correction, or omit completely to have bustools create its own “on list” for correction
- <sum>: What additional matrix to create by adding up the output matrices (options: cell, nucleus, or total)

Note: In the workflow above, we can additionally set the following two options in kb count (otherwise, the defaults are chosen):

- **--parity=single or --parity=paired**
- **--strand=forward** or **--strand=reverse** or **--strand=unstranded**

One can alternatively set those options at the end of <tech>, e.g.: <tech>%forward%paired

In addition to single-cell and single-nucleus RNA-seq, kb count can be used for bulk RNA-seq. Bulk RNA-seq generally does not have UMIs or cell barcodes (although artificial unique sample-specific barcodes are used to identify each sample) and relies on coding DNA mapping. With -x BULK as the technology string, a workflow specific for bulk RNA-seq quantification is executed (**Box 6**). This will produce both transcript-level and gene-level abundances that can be used by DESeq2^47,48^, sleuth^49^, limma-voom^50,51^, and other differential gene expression programs.

#### Box 6: kb count: bulk RNA-seq

Below, we show how to run kb count for preprocessing bulk RNA-seq data (**Box 4**). The procedure is similar to the preprocessing of single-cell RNA-seq, but there are some differences in how quantification is performed and barcode error correction is not performed due to the lack of cell barcodes in bulk RNA-seq. **--tcc** specifies that estimated counts should be produced in accordance with the count estimation algorithm in the original kallisto publication and **--matrix-to-directories** means that those quantifications should be reformatted into directories of “abundance files” with each sample being a different directory. The abundance files can be directly used by downstream tools designed for bulk RNA-seq differential gene expression. Below is an example usage for a paired-end unstranded bulk RNA-seq experiment on one sample.

**kb count -x BULK -o output_dir -<u>i index.idx</u> -g <u>t2g.txt</u> \**

**--parity=paired --strand=unstranded \**

**--tcc --matrix-to-directories <u>R1.fastq R2.fastq</u>**

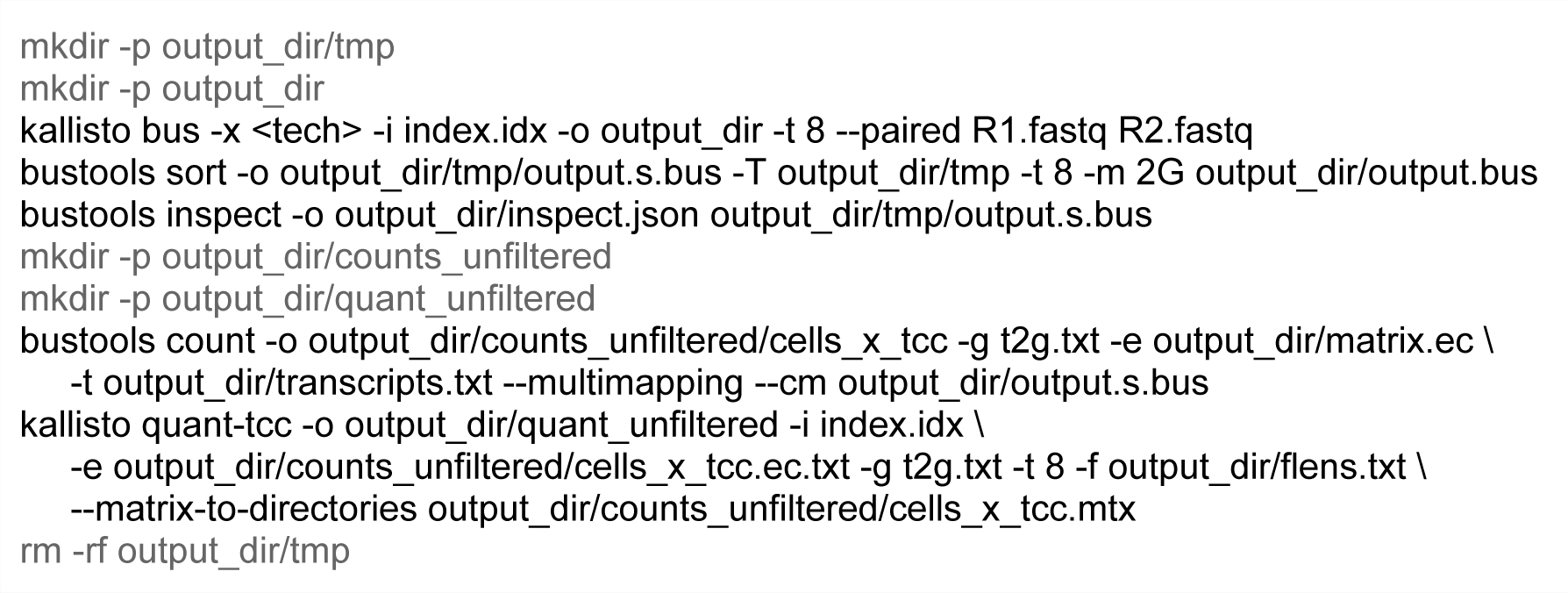

To facilitate multi-sample analysis, artificial unique sample-specific barcodes can be created and stored in the BUS file and the resulting mapping between the artificially generated barcode and the sample ID is outputted. These sample-specific barcodes are 16-bp in length and are also stored in the BUS file. Where there exists both a cell barcode (like in single-cell RNA-seq) and a sample-specific barcode, both sets of barcodes will be outputted so that each entry in the resulting output count matrix can be associated with a particular cell and a particular sample. To utilize the multi-sample workflow, a batch file containing the file names of the FASTQ files must be provided (**Box 7**).

#### Box 7: kb count (multi-sample analysis using the standard index type)

Below, we show how to run kb count to perform an analysis of multiple samples using the standard (default) index type. Use of this index type facilitates a workflow that is similar to the single-sample standard workflow (**Box 4**). A batch file (**batch.txt**) should be provided, in lieu of FASTQ files, listing all the samples to be analyzed with the paths to their respective FASTQ files. The **--batch-barcodes** option is provided in order to store the sample-specific barcodes that are created in addition to the cell barcodes (without this option, only cell barcodes are stored). This option can be omitted in the case that no cell barcodes exist (as in bulk RNA-seq).

**kb count -x <tech> -w <u>onlist.txt</u> -o output_dir -i <u>index.idx</u> -g <u>t2g.txt</u> \**

**--batch-barcodes <u>batch.txt</u>**

The only difference in the underlying kallisto command is in the *kallisto bus* command.

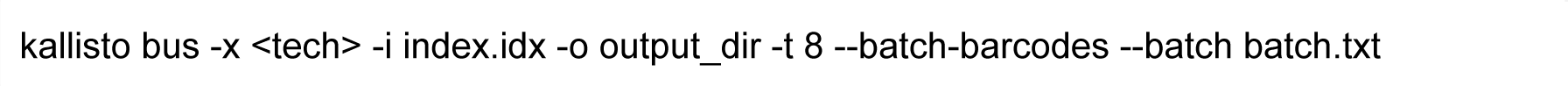

The batch.txt file looks as follows:

batch.txt

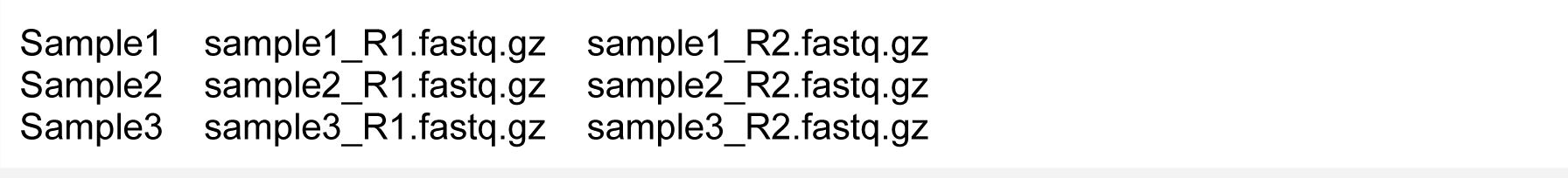

The sample ID is in the first column. Multiple rows can be provided for the same sample ID (e.g. if the FASTQ files are divided across multiple lanes). The third column can be omitted if only one FASTQ file is specified by the technology.

In the output directory (output_dir), there will be two files: matrix.cells (which contains the sample ID) and matrix.sample.barcodes (which contains the 16-bp sample-specific barcodes). Each line in matrix.cells corresponds to the same line in matrix.sample.barcodes. In the example above, the files look as follows:

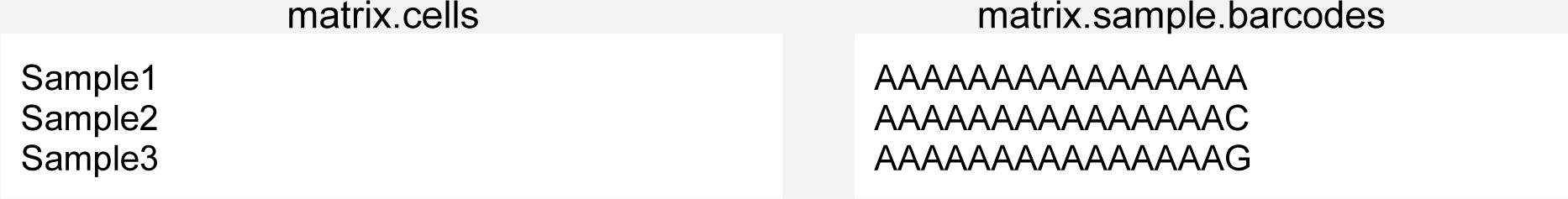

The technical details of how kb count utilizes kallisto and bustools are detailed in the following paragraph. Note that the **--dry-run** option in kb count outputs the kallisto and bustools commands that will be run without actually running the programs. Also, the **--verbose** option in kb count is helpful for examining the kallisto and bustools commands that are being run as well as their output.

kb count first invokes the **kallisto bus** command within kallisto to produce a BUS file, which stores the read mapping information, and then uses bustools^22^ commands to process the BUS file. The **kallisto bus** command maps RNA-seq reads to a kallisto index, and the resultant BUS file stores the mapping information, including the barcode, unique molecular identifier (UMI), and the equivalence class representing the set of transcripts the read is compatible with.^22^ In certain RNA-seq assays, barcodes and/or UMIs may not be present, and are therefore not considered when processing the BUS file. After the mapping step is complete, the BUS file is sorted via the **bustools sort** command to facilitate further processing. For single-cell and single-nucleus experiments with multiplexed barcodes in the sequencing reads, an “on list” of barcodes, representing the known barcode sequences that are included in the assay, needs to be provided. If an “on list” is unavailable, the **bustools allowlist** command can be used to construct one from a sorted BUS file. The barcodes in the sorted BUS file are error-corrected to the “on list” via **bustools correct**, then the BUS file is sorted again with **bustools sort**. The final sorted, on list-corrected BUS file is then used to generate quantifications via count matrices through the **bustools count** command. At any point, a sorted BUS file can be inputted into **bustools compress** to create a compressed BUS file (a BUSZ file), which can be subsequently decompressed via **bustools decompress**.^52^ Other bustools features enable more specialized workflows beyond what is provided by kb-python (**Supplementary Manual**).^1,53^

Quantification of RNA species can be performed in multiple ways as follows:

- Gene-level count matrices: In single-cell and single-nucleus RNA-seq, typically a gene-level count matrix is produced by collapsing UMIs to the gene level. Here, the bustools count command is run with the **--genecounts** option supplied. The **--umi-gene** option may also be provided for sequencing technologies where the UMIs are not expected to be unique within each cell. This ensures that in a case where two reads with the same UMI sequence map to different genes, they are considered to be two distinct molecules which were unintentionally labeled with the same UMI, and hence each gene gets a count. Such instances occur very frequently when UMIs are short such as in CEL-Seq2.^54^ By default, UMIs assigned to multiple genes after collapsing are discarded in quantification; however, the **--multimapping** option retains such UMIs and distributes the count uniformly across the assigned genes. This option, while improving the sensitivity of gene detection, causes non-integer counts to be created and is therefore disabled by default, consistent with other single-cell RNA-seq software. Finally, If one wishes to not perform UMI collapsing (i.e. each mapped read is its own unique molecule regardless of the UMI sequence), one can supply the **--cm** option for quantification.
- Transcript-level count matrices: Transcript-compatibility counts (TCCs) are counts assigned to equivalence classes where each equivalence class is defined by a unique set of transcripts. For producing a matrix of transcript-compatibility counts (TCCs), the **--genecounts** option is *not* provided, and **--multimapping** *is* provided to avoid discarding reads or collapsed UMIs that are assigned to multiple genes. If UMIs are not present in the sequencing technology, the **--cm** option is supplied to perform counting without UMI collapsing. While downstream analyses can be performed on TCCs^55,56^, it is more often useful to produce transcript-level abundances from the TCCs for technologies where sequencing reads span the full length of transcripts, such as bulk RNA-seq. In such cases, an expectation-maximization algorithm is typically performed to probabilistically estimate transcript abundances.^14,57^ The procedure to generate transcript-level abundance matrices is performed by running the **kallisto quant-tcc** command on the TCC matrices.

### Interfacing with genomic data specification and querying tools

seqspec^40^ provides a specification for the structure of genomic sequencing assays, formatted in a machine-readable YAML file. The specification can be readily inputted into the kallisto bustools workflow for preprocessing reads from a given assay (**Box 8**).

#### Box 8: Using seqspec specifications

**kb count -i index.idx -g t2g.txt \**

**-x $(seqspec index -t kb -m RNA -r R1.fastq,R2.fastq spec.yaml) \**

**-o output_dir -w $(seqspec onlist -m RNA -r barcode spec.yaml) R1.fastq R2.fastq**

Here, $() is process substitution.

This protocol as a whole can be executed on publicly available sequencing data using as few as two commands in addition to the installation command. This is made possible by genomic data and metadata command-line querying tools. Although many such tools exist, here, we utilize gget ref^58^, which can fetch reference genome FASTA files and genome annotation GTF files, and ffq^59^, which fetches the URL of the sequencing reads based on metadata retrieval (**Box 9**).

#### Box 9: Three commands to preprocess public RNA-seq reads with gget and ffq

Example with a mouse (*Mus musculus*) tissue single-cell RNA sequencing sample (SRA accession ID: SRR17068590), prepared using 10x (version 3) chemistry.

**pip install kb-python gget ffq**

**kb ref -i index.idx -g t2g.txt -f1 cdna.fasta $(gget ref --ftp -w dna,gtf mus_musculus)**

**kb count -i index.idx -g t2g.txt -x 10xv3 -o output_dir \**

**$(ffq --ftp SRR17068590 | jq -r ’.[] | .url’ | tr ’\n’ ’ ’)**

Notes: gget version 0.27.9, ffq version 0.3.0

## Anticipated Results

Here, the quantification output of the kb count command is described. While the initial step of kb count uses kallisto to produce a BUS file located at output_dir/output.bus, the actual quantification results are located in matrices in subdirectories of output_dir/. All matrices have the extension .mtx and will be in a sparse matrix (Matrix Market) file format with the barcodes (i.e. the cells or samples) being the matrix rows and the genes (or transcripts or equivalence classes or other features^60^) being the matrix columns.

### Gene-level counting

Gene-level counting to produce gene count matrices is the most common form of quantification for UMI-based single-cell and single-nucleus RNA-seq assays.

The **output_dir/counts_unfiltered/** directory contains the following information for <u>gene count matrices</u> (these are the matrices that are most commonly used for single-cell and single-nucleus RNA-seq analysis):

- <u>standard index type</u>

○ **cells_x_genes.mtx**: The count <u>matrix</u> (in Matrix Market file format); only exonic reads are counted
○ **cells_x_genes.barcodes.txt**: The barcodes representing the matrix <u>row names</u>
○ **cells_x_genes.genes.txt**: The gene IDs representing the matrix <u>column names</u>
○ **cells_x_genes.genes.names.txt**: Same as cells_x_genes.mtx except with gene names instead of gene IDs for the matrix columns
○ **cells_x_genes.barcodes.prefix.txt**: If sample-specific barcodes are generated in addition to cell barcodes being recorded, then this file will be created and the sample-specific barcodes will be stored here. The lines of this file correspond to the lines in the cells_x_genes.barcodes.txt which contains the cell barcodes (both files will have the same number of lines). The sample-specific barcodes and cell barcodes can be joined together as a unique identifier for downstream analysis.
- <u>nac index type</u>: same as the standard index type except the .mtx files produced are different

○ **cells_x_genes.mature.mtx**: The <u>mature RNA</u> count matrix
○ **cells_x_genes.ambiguous.mtx**: The <u>nascent RNA</u> count matrix
○ **cells_x_genes.nascent.mtx**: The <u>ambiguous RNA</u> count matrix
○ **cells_x_genes.cell.mtx**: The mature+ambiguous RNA count matrix (note: this is what is quantified in the count matrix with the standard index type workflow option)
○ **cells_x_genes.nucleus.mtx**: The <u>nascent+ambiguous RNA</u> count matrix
○ **cells_x_genes.total.mtx**: The <u>mature+nascent+ambiguous RNA</u> count matrix

### Transcript-level counting

For RNA-seq assays (e.g. bulk RNA-seq or Smartseq2) that profile the full length of transcripts in which case it is desirable to perform transcript-level quantification, the **--tcc** option is used.

The first step to doing transcript-level quantification is to obtain transcript-compatibility counts (TCCs) over equivalence classes (ECs). The TCCs will be outputted into **output_dir/counts_unfiltered/** which contains the following files for the standard workflow:

- **cells_x_tcc.mtx**: The count <u>matrix</u> containing the TCCs
- **cells_x_tcc.barcodes.txt**: The barcodes representing the matrix <u>row names</u>
- **cells_x_tcc.ec.txt**: The equivalence classes representing the matrix <u>column names</u> (note: this file has two columns – the first is the equivalence class numbers, which represent the column names, and the second is a comma-separated list of transcript numbers (0 based) for all transcripts within the equivalence class)

The --tcc option will additionally produce transcript-level estimated counts which will be placed in the **output_dir/quant_unfiltered/** directory which contains the following:

- **matrix.abundance.mtx**: The <u>matrix</u> containing the transcript-level estimated counts
- **matrix.abundance.tpm.mtx**: The <u>matrix</u> containing the TPM-normalized transcript-level abundances
- **matrix.efflens.mtx**: A <u>matrix</u> that contains the transcript effective lengths matrix.fld.tsv: A file with two columns, containing the mean and standard deviation, respectively, of the fragment length distribution used to produce transcript-level abundances and effective lengths for each row of the matrix.
- **matrix.abundance.gene.mtx**: A <u>matrix</u> that is the same as the matrix.abundance.mtx matrix except counts are aggregated to gene-level
- **matrix.abundance.gene.tpm.mtx**: A <u>matrix</u> that is the same as the matrix.abundance.tpm.mtx matrix except TPMs are aggregated to gene-level
- **transcripts.txt**: The transcript names representing the matrix <u>column names</u> for the transcript-level quantification matrices
- **genes.txt**: The gene IDs representing the matrix <u>column names</u> for the gene-level aggregation quantification matrices
- **transcript_lengths.txt**: The transcript names along with their lengths

*Note: The row names are the individual samples and will be the same as those in output_dir/counts_unfiltered/cells_x_tcc.barcodes.txt - The output_dir/matrix.cells and output_dir/matrix.sample.barcodes files provide a mapping between the sample name and the sample barcode.

*Note: The --matrix-to-directories option will output each row of the matrix into a separate subdirectory. In other words, using this option will produce multiple new directories within output_dir/quant_unfiltered/. Each one will be named abundance_{n} (where {n} is the sample number, corresponding to the rows in the matrix files). Within each subdirectory, an abundance.tsv text file and abundance.h5 HDF5 file will be created containing the quantifications for that particular sample. These abundance files are identical to the abundance files produced by the original version of kallisto for bulk RNA-seq.

### Loading single-cell output into downstream tools

To load the quantification results into SCANPY^61^ for downstream processing in python, an anndata^62^ object needs to be created. A user can import the count matrices into an anndata object (**Box 10**), or can run kb count with the --h5ad option to generate the anndata object directly. A user can also create a loom file directly by running kb count with the --loom option.

#### Box 10: Loading count matrices into scanpy

The standard index type produces a single count matrix (in output_dir/counts_unfiltered/) which can be loaded into scanpy via an anndata object as follows:

**import kb_python.utils as kb_utils**

**adata = kb_utils.import_matrix_as_anndata(“cells_x_genes.mtx”,**

**“cells_x_genes.barcodes.txt”,**

**“cells_x_genes.genes.names.txt”)**

The nac index type produces multiple count matrices. If one wishes to investigate different RNA species separately, one can load multiple count matrices as *layers* into the anndata object. The first layer will always be named “spliced” and the second layer will always be named “unspliced”. Below, we load in the “spliced” layer (from the cells_x_genes.cell.mtx count matrix which represents mature+ambiguous counts) and the “unspliced” layer (from the cells_x_genes.nascent.mtx count matrix, which represents the nascent counts).

**import kb_python.utils as kb_utils**

**adata_spliced = kb_utils.import_matrix_as_anndata(“cells_x_genes.cell.mtx”,**

**“cells_x_genes.barcodes.txt”,**

**“cells_x_genes.genes.names.txt”)**

**adata_unspliced = kb_utils.import_matrix_as_anndata(“cells_x_genes.nascent.mtx”,**

**“cells_x_genes.barcodes.txt”,**

**“cells_x_genes.genes.names.txt”)**

**adata = kb_utils.overlay_anndatas(adata_spliced, adata_unspliced)**

Note: If sample-specific barcodes are specified in addition to cell barcodes, one can add `batch_barcodes_path=“cells_x_genes.barcodes.prefix.txt”` to import_matrix_as_anndata to concatenate the two barcodes together.

If one wishes to write an anndata object to a loom file, one can simply do:

**adata.write_loom(“/path/to/loom/file.loom”)**

If one runs kb count with the --h5ad or --loom option, the files adata.h5ad or adata.loom are created alongside the count matrix files. With the nac index type, the automatically generated adata.h5ad or adata.loom file will have three layers: nascent, mature, and ambiguous, containing those respective count matrices. For the adata.h5ad file, one can read it in by the following:

**adata = anndata.read_h5ad(“output_dir/counts_unfiltered/adata.h5ad”)**

Note: anndata version 0.9.2; python version 3.8.0

For downstream processing in R, one can load the quantification results into Seurat^63^ (**Box 11**). Additionally, in R, one can create a Bioconductor SingleCellExperiment^64^ object for use with single-cell analysis R packages such as scran^65^ and scater^66^ (**Box 12**).

#### Box 11: Loading count matrices into Seurat

The standard workflow produces a single count matrix (in output_dir/counts_unfiltered/), which can be loaded into Seurat as follows:

**library(Seurat)**

**expression_matrix <-ReadMtx(mtx=“cells_x_genes.mtx”,**

**features = “cells_x_genes.genes.names.txt”,**

**cells = “cells_x_genes.barcodes.txt”,**

**feature.column=1,**

**mtx.transpose = TRUE)**

For single-nucleus RNA-seq, one would use the nac workflow and the count matrix that should be loaded in would be cells_x_genes.nucleus.mtx (or cells_x_genes.total.mtx if one wishes to add in the mature RNA counts).

Notes: Seurat version 4.0.5; R version 4.1.1

#### Box 12: Loading count matrices into SingleCellExperiment

Here, we show how to build a SingleCellExperiment object in R from the standard workflow output count matrix (in output_dir/counts_unfiltered/):

**library(SingleCellExperiment)**

**library(Matrix)**

**counts <-Matrix::readMM(“cells_x_genes.mtx”)**

**gene_ids <-readLines(“cells_x_genes.genes.txt”)**

**gene_symbols <-readLines(“cells_x_genes.genes.names.txt”)**

**barcodes <-readLines(“cells_x_genes.barcodes.txt”)**

**sce `<-` SingleCellExperiment(list(counts=t(counts)),**

**colData=DataFrame(Barcode=barcodes),**

**rowData=DataFrame(ID=gene_ids,SYMBOL=gene_symbols))**

**rownames(sce) `<-` gene_ids**

Notes: SingleCellExperiment version 4.0.5; R version 4.1.1

The count matrices are initially unfiltered, which makes them very large and inefficient to process. After filtering for cells with sufficient UMI counts (among other criteria), the matrices that are loaded in will become much smaller and more efficient to process.

## Materials

- A 64-bit computer running either macOS, Windows, or a Linux/Unix operating system.
- kb-python version 0.28.2 or later

○ kallisto version 0.50.1 or later (which comes packaged with kb-python)
○ bustools version 0.43.2 or later (which comes packaged with kb-python)
- Python 3.7 or later (for kb-python version 0.28.2)
- Bulk, single-cell, or single-nucleus RNA sequencing reads in (possibly gzip) FASTQ format.

### Timing

The runtime depends on the size of the reference being indexed, the number and length of the sequencing reads being processed, other properties of the dataset being quantified, system hardware, and the number of threads allotted. The kb ref command only needs to be run once to create the index against which reads will be mapped. With 8 threads on a server with x86-64 architecture and 32 Intel Xeon CPUs (E5-2667 v3 @ 3.20GHz), kb ref, which by default uses the d-list option, takes approximately 15 minutes to generate a standard index from the GRCm39 mouse genome (using the respective raw *unfiltered* GTF file) and an hour to generate the nac index. For the preprocessing of 800 million Illumina sequencing reads (stored in a single pair of fastq.gz files) produced by single-cell RNA-seq from 10x Genomics, kb count with the nac workflow can take under an hour on 8 threads and under 40 minutes on 16 threads, with an even lower runtime for the standard workflow.

### Troubleshooting

The **--verbose** option in kb ref and kb count is helpful for examining the kallisto and bustools commands that are being run as well as their output. This can be used to troubleshoot errors.

The **--overwrite** option in kb ref and kb count can be used to regenerate output files and directories that were produced from a previous kb-python run.

The output directory of a kb count run contains multiple JSON^67^ files that contain quality control values such as the percentage of reads pseudoaligned.

If one receives an “Error: incompatible indices”, either the index file being supplied is corrupted, is not an actual kallisto index file, or was an index file generated by a version of kallisto that utilized a different index format; kallisto version 0.50.1 utilizes a different index format than previous versions and future versions of kallisto may likely adopt a newer index format. In any case the index should be regenerated.

The t2g (transcripts-to-gene mapping) file created by kb ref should be the exact file used by kb count when running kb count on that index. All the transcripts in the t2g file must be exactly the same as the transcripts present in the kallisto index. Incompatibilities can lead to unpredictable behavior in the bustools quantification step.

When using kb ref to generate a kallisto index, a genome FASTA file (not a transcriptome FASTA file) should be supplied along with the genome annotation GTF file. A transcriptome file will automatically be generated by kb ref and be indexed by kallisto. In general, the Ensembl^30^ .dna.toplevel.fa.gz files or the GENCODE^68^ .primary_assembly.genome.fa.gz files should be used as the reference genome. Use of FASTA files incompatible with the supplied GTF will lead to errors.

When performing multiple kb-python runs simultaneously, a different temporary directory must be specified via the --tmp option for each run. The temporary directory also must not exist beforehand.

Finally, one should make sure that the value supplied to the -x technology string option matches the assay from which the sequencing reads were generated.

> Note: If the technology string begins with a -, for example: -1,0,0:0,0,5:0,5,0, one would need to write -x “ -1,0,0:0,0,5:0,5,0” to avoid the string being misinterpreted as a command-line flag.

### Procedure

Here, we describe the procedures to use for mouse samples of paired-end bulk RNA-seq, 10x (version 3) single-cell RNA-seq, and 10x (version 3) single-nucleus RNA-seq.

#### Bulk RNA-seq

**Input:**

- Paired-end unstranded mouse RNA-seq reads (3 samples): sample1_R1.fastq.gz sample1_R2.fastq.gz sample2_R1.fastq.gz sample2_R2.fastq.gz sample3_R1.fastq.gz sample3_R2.fastq.gz

1. **Install kb-python**

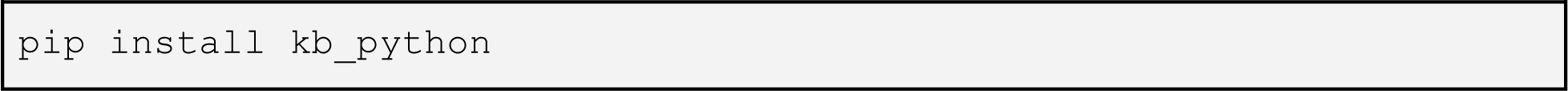
2. **Download the mouse genome and annotation files**

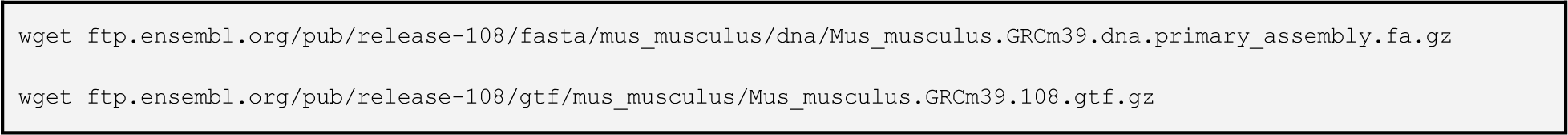
3. **Build the index**

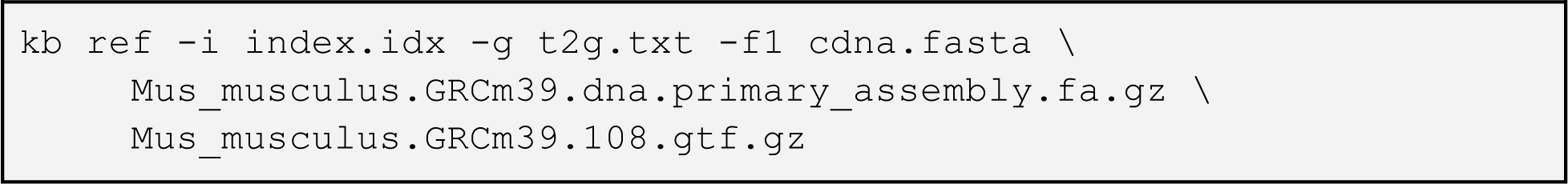
4. **Map the input sequencing reads to the index**

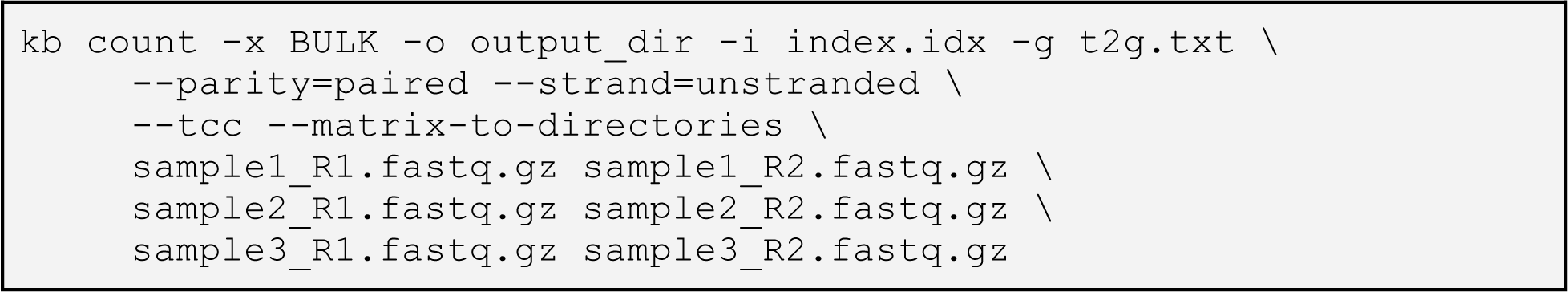
5. **Analyze the output**

Output for sample 1:

- output_dir/quant_unfiltered/abundance_1/abundance.tsv
- output_dir/quant_unfiltered/abundance_1/abundance.gene.tsv
- output_dir/quant_unfiltered/abundance_1/abundance.h5

Output for sample 2:

- output_dir/quant_unfiltered/abundance_2/abundance.tsv
- output_dir/quant_unfiltered/abundance_2/abundance.gene.tsv
- output_dir/quant_unfiltered/abundance_2/abundance.h5

Output for sample 3:

- output_dir/quant_unfiltered/abundance_3/abundance.tsv
- output_dir/quant_unfiltered/abundance_3/abundance.gene.tsv
- output_dir/quant_unfiltered/abundance_3/abundance.h5

The abundance.tsv files contain the transcript-level abundances. The abundance.h5 file contains the same information as the abundance.tsv files except in HDF5 format. The abundance.gene.tsv files contain the gene-level abundances (taken by summing up the transcript-level abundances for each gene). These files can be used in downstream differential gene expression programs.

#### Single-cell RNA-seq

**Input:**

- 10x version 3 single-cell RNA-seq reads: R1.fastq.gz and R2.fastq.gz

1. **Install kb-python**

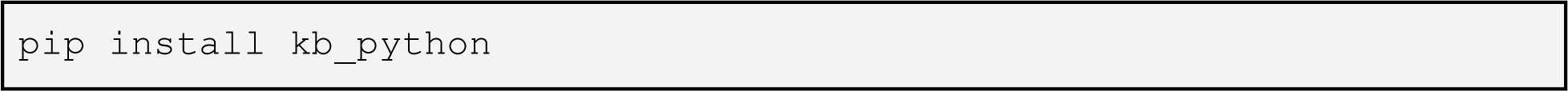
2. **Download the mouse genome and annotation files**

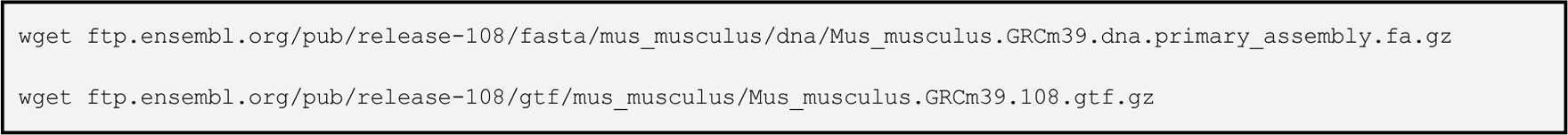
3. **Build the index**

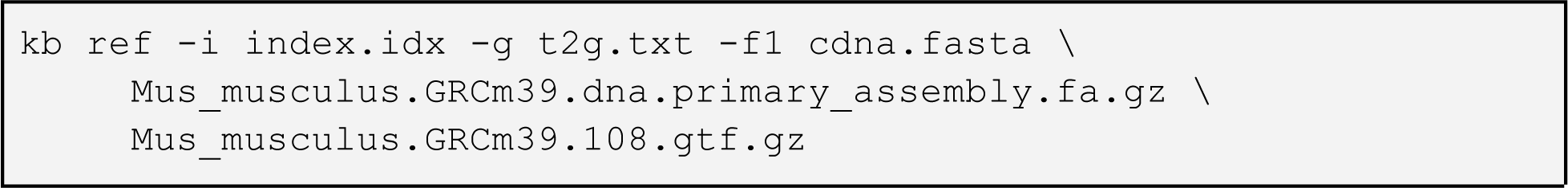
4. **Map the input sequencing reads to the index**

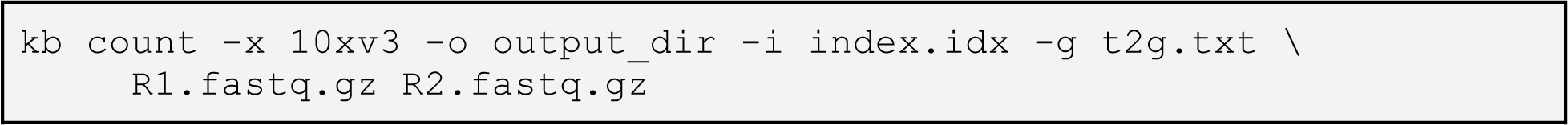
5. **Analyze the output**

Output:

- output_dir/counts_unfiltered/cells_x_genes.mtx
- output_dir/counts_unfiltered/cells_x_genes.barcodes.txt
- output_dir/counts_unfiltered/cells_x_genes.genes.txt
- output_dir/counts_unfiltered/cells_x_genes.genes.names.txt

The cells_x_genes.mtx is the count matrix file with the barcodes (the row names) listed in cells_x_genes.barcodes.txt and the gene names (the column names) listed in cells_x_genes.genes.names.txt (for gene IDs instead of gene names, use cells_x_genes.genes.txt).

## Single-nucleus RNA-seq

**Input:**

- 0x version 3 single-nucleus RNA-seq reads: R1.fastq.gz and R2.fastq.gz

1. **Install kb-python**

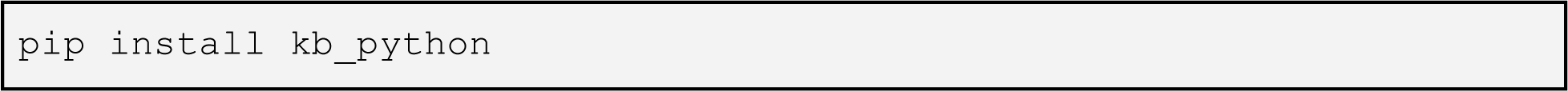
2. **Download the mouse genome and annotation files**

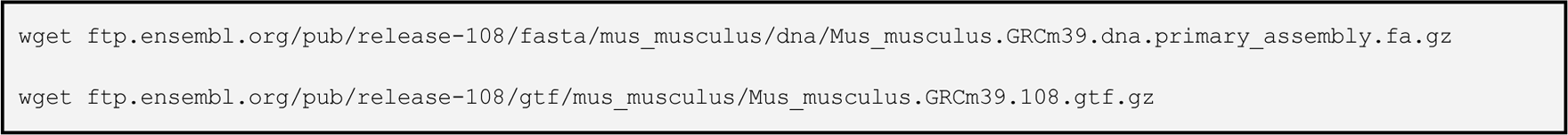
3. **Build the index**

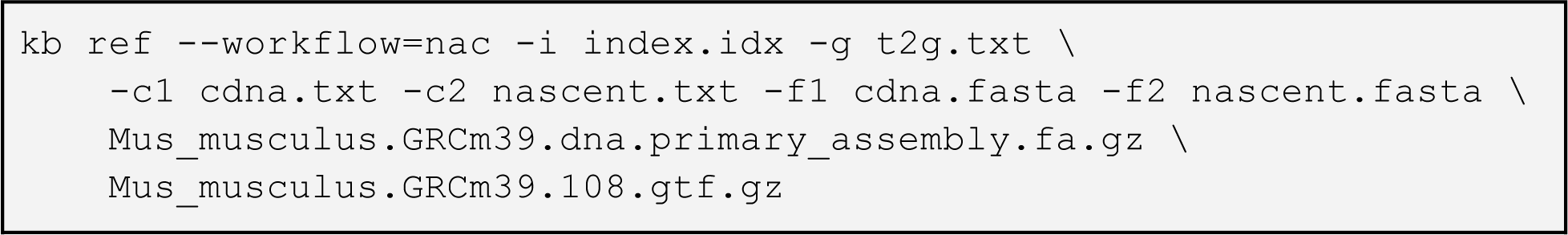
4. **Map the input sequencing reads to the index**

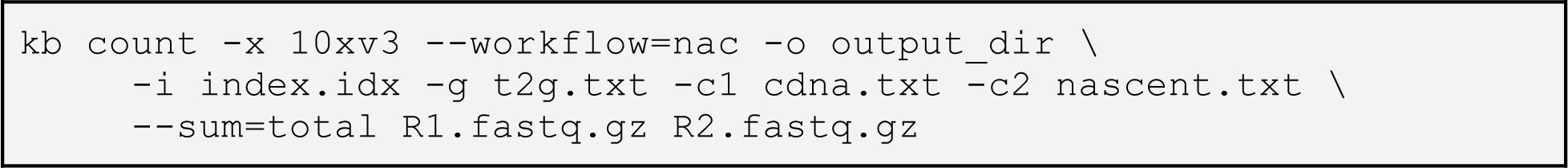
5. **Analyze the output**

Output:

- output_dir/counts_unfiltered/cells_x_genes.mature.mtx
- output_dir/counts_unfiltered/cells_x_genes.nascent.mtx
- output_dir/counts_unfiltered/cells_x_genes.ambiguous.mtx
- output_dir/counts_unfiltered/cells_x_genes.cell.mtx
- output_dir/counts_unfiltered/cells_x_genes.nucleus.mtx
- output_dir/counts_unfiltered/cells_x_genes.total.mtx
- output_dir/counts_unfiltered/cells_x_genes.barcodes.txt
- output_dir/counts_unfiltered/cells_x_genes.genes.txt
- output_dir/counts_unfiltered/cells_x_genes.genes.names.txt

This workflow can be used for both single-cell RNA-seq and single-nucleus RNA-seq. Many count matrix files (.mtx files) are generated. For quantification of total RNA present in each cell or nucleus, one would want to use the cells_x_genes.total.mtx. For biophysical models that jointly consider spliced and unspliced transcripts, one may want to use cells_x_genes.cell.mtx (for the “spliced” transcripts) and cells_x_genes.nascent.mtx (for the “unspliced” transcripts).

The barcodes (the matrix row names) are listed in cells_x_genes.barcodes.txt and the gene names (the matrix column names) are listed in cells_x_genes.genes.names.txt (for gene IDs instead of gene names, use cells_x_genes.genes.txt).

### Additional extensions

There are many ways to extend the standard workflows beyond bulk RNA-seq, 10x single-cell RNA-seq, and 10x single-nucleus RNA-seq. For an additional, extended example that involves preprocessing mouse multiplexed single-nucleus SPLiT-seq RNA-seq data with a filtered mouse genome annotation, see **Supplementary Tutorial**.

## Contributions

All authors contributed either directly to kallisto, bustools, kb-python or to the methods implemented in the software. D.K.S. led the development of the latest versions (at the time of writing this manuscript) of kallisto (version 0.50.1), bustools (version 0.43.2), and kb-python (version 0.28.2). N.L.B. conceived kallisto. A.S.B. conceived kb-python and K.M.H. created, implemented, and developed kb-python under the supervision of A.S.B. L.M. identified the need for creating a transcriptome FASTA file coherent with a genome and GTF as implemented in kb-python. A.S.B and P.M. implemented the initial version of bustools and its interface with kallisto, which was published in Melsted, Booeshaghi et al., 2021^1^ where early versions of these software were benchmarked. D.K.S. and K.E.H. implemented the d-list option in kallisto and adapted kallisto to use the Bifrost de Bruijn graph with help from G.H. L.L. conceived the translated search (the --aa option)^69^ and implemented it with help from D.K.S. and K.E.H. J.G. augmented the functionalities of bustools. D.K.S. refactored kallisto providing additional modularity with respect to the expectation-maximization algorithm implemented by H.P., and unifying the treatment of bulk and single-cell data. P.M. and L.P. supervised the initial development and coordination of kallisto and bustools. D.K.S. drafted the initial manuscript. All authors edited and reviewed the final manuscript.

## Competing interests

The authors declare no competing financial interests.

## Key reference using this protocol

Melsted, P., Booeshaghi, A.S. *et al.* Modular, efficient and constant-memory single-cell RNA-seq preprocessing. *Nat Biotechnol* 39, 813–818 (2021). https://doi.org/10.1038/s41587-021-00870-2

## Acknowledgements

D.K.S. was funded by the UCLA-Caltech Medical Scientist Training Program (NIH NIGMS training grant T32 GM008042). L.P. was supported in part by the National Institutes of Health (NIH) grants U19MH114830 and 5UM1HG012077-02. A.S.B. was funded in part by Icelandic Research Fund Project grant number 218111-051. Other contributors to the software and methods include Vasilis Ntranos, Lauren Liu, Fan Gao, Eduardo da Veiga Beltrame, and Jase Gehring. The development of kallisto and bustools was also funded in part by a grant awarded during round 2 of the Essential Open Source Software for Science by the Chan Zuckerberg Initiative for “Open Source Software for Bulk and Single-cell RNA-seq”.

## Code Availability

The kallisto software is available at https://github.com/pachterlab/kallisto. The bustools software is available at https://github.com/BUStools/bustools. The kb-python software is available at https://github.com/pachterlab/kb_python.

## Notes

### Competing Interest Statement

The authors have declared no competing interest.

### Summary of Updates

Clarified certain aspects of the protocol (when and how to use certain parameters and how to store and load anndata objects in python).

