## Supplementary Material for "kallisto, bustools, and kb-python for quantifying bulk, single-cell, and single-nucleus RNA-seq"

**Supplementary Note 1**: Installation of kallisto and bustools from source or installation of specific versions of the software.

##### Installing kallisto and bustools from source

kallisto (version 0.50.1):

```
git clone --branch v0.50.1 https://github.com/pachterlab/kallisto
cd kallisto
mkdir build
cd build
cmake ..
make
make install
```

bustools (version 0.43.1):

```
git clone --branch v0.43.1 https://github.com/BUStools/bustools
cd bustools
mkdir build
cd build
cmake ..
make
make install
```

Note: The --branch argument can be omitted to install the latest version of the software.

##### Using kb_python with kallisto and bustools installed from source

kb_python can be run with compiled binaries by supplying the paths to the binaries as follows:

```
kb ref --kallisto=/path/to/kallisto --bustools=/path/to/bustools …
```

```
kb count --kallisto=/path/to/kallisto --bustools=/path/to/bustools …
```

##### Installing a specific version of kb_python

A specific version of kb_python (e.g. version 0.28.0) can installed as follows:

```
pip install kb_python==0.28.0
```

**Supplementary Note 2**: Indexing a custom set of k-mers.

##### Indexing a custom set of k-mers

When multiple sequences may belong to the same “target”, as is the case with genetic polymorphisms, it can be desirable to index k-mers distributed across multiple targets rather than across a single contiguous target sequence. The target names in the input FASTA file must be numbers (specifically, zero-indexed numerical identifiers). Each k-mer in the target sequence is associated with the target name specified in the header line. Indexing this FASTA file can then be accomplished in the custom workflow using the **--distinguish** keyword.

**custom** workflow (--distinguish):

```
kb ref --workflow=custom -i index.idx --distinguish custom.fasta
```

```
kallisto index -t 8 -i index.idx --distinguish custom.fasta
```

Example custom.fasta file (with 3 targets):

```
>0
ACTCTATCATCATCTACTACTACTCGCAGCGACGACATCAGCTTTTTT
>1
GCGCGCCGCCGACGACACGCAGAGAAGAAAGCGCGACGAC
>2
TTATGTGTCGTGTAGTCGTAGTGTGTCGTGCCGCCGCGCGCAAA
>2
ATATACGATCATCAGCGACAGACTACTTCAGAAGACTATCA
>0
GTCGATCGGTGTCACATGCGCAAGCGTCAGCGACACGACTTCGG
```

##### D-listing a custom set of k-mers

When FASTA sequences are supplied to **--d-list**, distinguishing flanking k-mers (DFKs) are extracted from those sequences and placed in a D-list. Reads containing D-list k-mers will not be mapped. One can also specify a custom set of k-mers to be in the D-list, by using an empty sequence header. In the following example, since the header is absent, *all* k-mers in the sequence will be D-listed (if a header were present, only DFKs would be D-listed).

```
>
ACGCGACATAGCAGACTAGACATTATTTACGTATTATGATAGTAGAT
```

**Supplementary Note 3**: Filtering GTF entries when constructing the reference.

**kb ref: --include-attributes and --exclude-attributes to filter GTF entries**

Specific GTF entries can be included or excluded when building a reference transcriptome from a genome FASTA and GTF file. This can be done by using the following arguments to kb ref:

--include-attribute KEY:VALUE

--exclude-attribute KEY:VALUE

Where KEY is the name of the field (e.g. gene_biotype) in the GTF file and the VALUE is the value of the field (e.g. protein_coding).

The box below shows an example of how to use --include-attribute to include only certain gene biotypes (the remaining gene biotypes present in the GTF file will not be included). Note that these are the same biotypes included in the Ensembl GRCh38 Cell Ranger reference (as of Cell Ranger version 7.1.0).

```
kb ref -i index.idx -g t2g.txt -f1 cdna.fasta \
    --include-attribute gene_biotype:protein_coding \
    --include-attribute gene_biotype:lncRNA \
    --include-attribute gene_biotype:lincRNA \
    --include-attribute gene_biotype:antisense \
    --include-attribute gene_biotype:IG_LV_gene \
    --include-attribute gene_biotype:IG_V_gene \
    --include-attribute gene_biotype:IG_V_pseudogene \
    --include-attribute gene_biotype:IG_D_gene \
    --include-attribute gene_biotype:IG_J_gene \
    --include-attribute gene_biotype:IG_J_pseudogene \
    --include-attribute gene_biotype:IG_C_gene \
    --include-attribute gene_biotype:IG_C_pseudogene \
    --include-attribute gene_biotype:TR_V_gene \
    --include-attribute gene_biotype:TR_V_pseudogene \
    --include-attribute gene_biotype:TR_D_gene \
    --include-attribute gene_biotype:TR_J_gene \
    --include-attribute gene_biotype:TR_J_pseudogene \
    --include-attribute gene_biotype:TR_C_gene \
    genome.fasta genome.gtf
```

**Supplementary Manual**: Reference for kallisto and bustools commands.

### 1. kallisto

Running kallisto usually involves two steps: 1) Indexing a FASTA file of target sequences via `kallisto index`, and 2) Mapping sequencing reads to kallisto index using `kallisto bus`.

#### 1.1 kallisto index

Builds a kallisto index.

Usage: kallisto index [arguments] FASTA-files

```
Required argument:
-i, --index=STRING       Filename for the kallisto index to be constructed

Optional arguments:
-k, --kmer-size=INT      k-mer (odd) length (default: 31, max value: 31)

-t, --threads=INT        Number of threads to use (default: 1)

-d, --d-list=STRING      Path to a FASTA-file containing sequences to mask
                         from quantification (i.e. to extract distinguishing
                         flanking k-mers from).

--make-unique            Replace repeated target names with unique names

--aa                     Generate index from a FASTA-file containing
                         amino acid sequences

--distinguish            Generate index where sequences are distinguished
                         by the sequence name, for example, when indexing
                         k-mers distributed across multiple targets rather
                         than across a single contiguous target sequence.

-T, --tmp=STRING         Temporary directory (default: tmp)

-m, --min-size=INT       Length of minimizers (default: automatically chosen)

-e, --ec-max-size=INT    Maximum number of targets in an equivalence class
                         (default: no maximum)
```

Among the optional arguments in `kallisto index`, in a general use case, typically only `-i` (`--index`; to specify the name of the index output filename), `-t` (`--threads`; to specify the number of threads), and `-d` (`--d-list`; to specify the filename from which to extract distinguishing flanking k-mers) are used.

#### 1.2 kallisto bus

Generates a BUS file containing the results from mapping sequencing reads to a kallisto index.

Usage:
kallisto bus [arguments] FASTQ-files
kallisto bus [arguments] --batch=batch.txt

```
Required arguments:
-i, --index=STRING          Filename for the kallisto index to be used for
                            pseudoalignment

-o, --output-dir=STRING     Directory to write output to

-x, --technology=STRING     The "technology" string for the sequencing
                            technology used

Other arguments:
-l, --list                  List the technologies that are hard-coded into
                            kallisto so the name of the technology can
                            simply be supplied as the technology string

-B, --batch=FILE            Path to a batch file. The batch file is a text
                            file listing all the samples to be analyzed
                            with the paths to their respective FASTQ files.
                            If a batch file is supplied, then one shouldn't
                            supply FASTQ files on the command line.

-t, --threads=INT           Number of threads to use (default: 1)

-b, --bam                   Input file is a BAM file rather than a set of
                            FASTQ files. Note: This is a nonstandard
                            workflow. It is strongly recommended to supply
                            FASTQ files rather than use this option and not
                            all technologies are supported by this option.

-n, --num                   Output read number in flag column of BUS file
```

The read number is zero-indexed. One can view the read numbers by inspecting the BUS file using *bustools text*. This option is useful for pulling specific mapped reads out of the FASTQ file or for examining which reads did not end up being mapped by kallisto. (Important note: BUS files with read numbers in the flag column can NOT be used in quantification tasks with bustools). (Note: incompatible with --bam)

-N, --numReads=INT

Maximum number of reads to process from supplied input. This is useful for processing a small subset of reads from a large sequencing experiment as a quick quality control. Moreover, the program returns 1 if the number of reads processed from the input is less than the number supplied here. This is useful for catching errors when we expect a certain number of reads to be present in the input but not all the reads end up being there.

-T, --tag=STRING

5′ tag sequence to identify UMI reads for certain technologies. This is useful for smart-seq3 where the UMI-containing reads have an 11-bp tag sequence (ATTGCGCAATG) located at the beginning of the UMI location. If this tag sequence is present immediately before the UMI location, then the UMI is processed into the output BUS file; for all other sequences, the UMI field in the BUS file is left empty (the field is populated with the value -1 in binary format).
Note: Matching the tag sequence is done with a hamming distance error tolerance of 1 if the tag is longer than 5 nucleotides. Otherwise, no error tolerance is permitted.
Note: If strand-specificity is enabled, it will only be applied to the UMI-containing reads.

--fr-stranded

Strand specific reads, first read forward

--rf-stranded

Strand specific reads, first read reverse

| | |
|---|---|
| `--unstranded` | Treat all read as non-strand-specific |
| `--paired` | Treat reads as paired (i.e. if two biological read sequences are present across two FASTQ files, they will be mapped taking into account their paired-endness: fragment length distribution will be estimated for the read pairs, and only one read in the pair needs to map successfully in order to be considered successful pseudoalignment) |
| `--aa` | Align to index generated from a FASTA-file containing amino acid sequences |
| `--inleaved` | Specifies that input is an interleaved FASTQ file. That is, only one FASTQ file is supplied and the sequences are interleaved. For example, instead of an R1 and R2 FASTQ file, a single FASTQ file can be supplied where the reads are listed in order of each R2 read immediately following each R1 read. This is also useful when piping interleaved output generated by another program directly into *kallisto bus* which can be done by supplying - as the input file in lieu of FASTQ file names. |
| `--batch-barcodes` | Records both the generated sample-specific barcodes as well as the cell barcodes extracted from the reads in the output BUS file. If not supplied, then the sample-specific barcodes are not recorded. |

In the output directory specified by `-o` or `--output-dir`, the following files are made:

- output.bus: A BUS file containing the mapped reads information, which will be further processed using bustools.
- transcripts.txt: A text file containing a list of the names of the targets or transcripts used.
- matrix.ec: A text file containing the equivalence classes. The equivalence class number (zero-indexed) is in the first column and a comma-separated list of target or transcript IDs belonging to that equivalence class are in the second column. The transcript IDs are numbers (zero-indexed) that correspond to the line numbers (zero-indexed) in the transcripts.txt file.
- run_info.json: Contains information about the run, including percent of reads

pseudoaligned, number of reads processed, index version, etc.
- flens.txt: Only produced when using paired-end mapping. Contains the fragment length distribution, which can be used by `kallisto quant-tcc` to produce TPM abundance values.

#### 1.3 kallisto quant-tcc

Quantifies abundance from pre-computed transcript-compatibility counts. It takes in a transcript compatibility counts (TCC) matrix outputted by `bustools count` and runs an expectation-maximization (EM) algorithm to produce transcript abundances. This is useful for producing TPM values from bulk RNA-seq and smart-seq2 RNA-seq data. The output files can be used by bulk RNA-seq differential gene expression programs.

Usage: kallisto quant-tcc [arguments] transcript-compatibility-counts-file

```
Required arguments:
-o, --output-dir=STRING       Directory to write output to


-e, --ec-file=FILE            File containing equivalence classes
                              (the equivalence class file in the same
                              directory as the output matrix file
                              should be used)

Other arguments:
-i, --index=STRING            Filename for the kallisto index
                              to be used
                              (required if --txnames is not supplied
                              or if any of the fragment length
                              options: -f, -l, -s, is supplied
                              since the index contains transcript
                              lengths, which is necessary for length
                              normalization)

-T, --txnames=STRING          File with names of transcripts
                              (required if index file not supplied)

-f, --fragment-file=FILE      File containing fragment length
                              distribution (flens.txt outputted by
                              kallisto)

-l, --fragment-length=DOUBLE  Estimated average fragment length
```

| | |
|---|---|
| -s, --sd=DOUBLE | Estimated standard deviation of fragment length<br>(note: -l, -s values only should be supplied when effective length normalization needs to be performed but --fragment-file is not specified)<br><br>Note: If none of the fragment length options: -f -l, -s, are supplied, then effective length normalization is not performed (i.e. transcript length isn't taken into account when quantification is performed). |
| -p, --priors=FILE | Priors for the EM algorithm, either as raw counts or as probabilities. Pseudocounts are added to raw counts to prevent zero valued priors. Supplied in the same order as the transcripts in the transcriptome (e.g. in --txnames). |
| -t, --threads=INT | Number of threads to use (default: 1) |
| -g, --genemap=FILE | File for mapping transcripts to genes (this is the t2g.txt file produced by kb ref in kb-python and is required for obtaining gene-level abundances) |
| -G, --gtf=FILE | GTF file for transcriptome information (can be used instead of --genemap for obtaining gene-level abundances) |
| -b, --bootstrap-samples=INT | Number of bootstrap samples (default: 0)<br>Bootstrap samples are useful for obtaining inferential variance which can be used by programs such as sleuth. |
| --matrix-to-files | Reorganize matrix output into abundance tsv files |
| --matrix-to-directories | Reorganize matrix output into abundance tsv files across multiple directories |

```
--seed=INT                 Seed for the bootstrap sampling
                           (default: 42)

--plaintext                Output plaintext only, not HDF5
                           (When --matrix-to-directories or
                           --matrix-to-files are supplied, HDF5
                           files are outputted by default, in
                           addition to the plaintext abundance tsv
                           files since HDF5 files containing
                           abundance information are used by
                           programs such as sleuth; this option
                           disables that).
```

In the output directory specified by `-o` or `--output-dir`, the following files are made:

- matrix.abundance.mtx: A sample-by-transcript (or cell-by-transcript) MatrixMarket sparse matrix file containing the estimated transcript counts.
- matrix.abundance.gene.mtx: A sample-by-gene (or cell-by-gene) MatrixMarket sparse matrix file containing the estimated transcript counts summed up to gene-level. Only made if a transcript-to-gene mapping was provided.
- matrix.abundance.tpm.mtx: A sample-by-transcript (or cell-by-transcript) MatrixMarket sparse matrix file containing the normalized transcript abundances (if effective length normalization is performed, then the results are in length-normalized TPM units; otherwise the results are in CPM [counts-per-million] units wherein each value is normalized by the sum of all counts for that particular sample or cell).
- matrix.abundance.gene.tpm.mtx: A sample-by-gene (or cell-by-gene) MatrixMarket sparse matrix file containing the same information as matrix.abundance.tpm.mtx except summed up to gene-level if a transcript-to-gene mapping was provided.
- transcripts.txt: A text file containing a list of the names of the targets or transcripts used (not made if a transcripts file was already provided via `--txnames`). These transcripts correspond to the columns of transcripts in the matrix abundance output files.
- genes.txt: A text file containing a list of genes, if a transcript-to-gene mapping was provided. These genes correspond to the columns of genes in the matrix abundance output files.
- --matrix-to-files: If this option is provided, the abundance output files will be named abundance_{n}.tsv and abundance_{n}.h5 (hdf5 format) where {n} is the sample number or cell number (which corresponds to the rows in the matrix files). If bootstrapping is enabled, additional abundance tsv files (starting with the prefix bs_abundance_{n}_) will be created for each bootstrap sample. If a transcript-to-gene mapping is provided, abundance.gene_{n}.tsv files will be created as well with the gene-level quantification.
- --matrix-to-directories: If this option is provided, directories named abundance_{n}

(where {n} is the sample number or cell number, corresponding to the rows in the matrix files) will be created. Within each directory, an abundance.tsv text file and abundance.h5 HDF5 file will be created containing the quantifications for that particular sample or cell. If bootstrapping is enabled, additional abundance tsv files (starting with the prefix bs_abundance_) will be created for each bootstrap sample. If a transcript-to-gene mapping is provided, an abundance.gene.tsv file will be created within each directory with the gene-level quantification.

The first few lines of an abundance tsv file looks as follows:

```
target_id            length   eff_length  est_counts    tpm
ENST00000641515.2    2618     2349.39     0              0
ENST00000426406.4    939      670.39      0              0
ENST00000332831.4    995      726.39      0              0
ENST00000616016.5    3465     3196.39     5.68407        0.128913
ENST00000618323.5    3468     3199.39     1.83535        0.041586
```

#### 1.3 kallisto quant

`kallisto quant` is an old usage of kallisto when kallisto was first developed for bulk RNA-seq quantification. It is now recommended that users use the `kallisto bus` command instead.

As such, documentation for the old `kallisto quant` is not within the scope of this protocol.

#### 1.4 kallisto inspect

Inspects and gives information about an index. The index can be loaded more quickly by using multiple threads, which can be specified by the **-t** option.

Example usage:

```
kallisto inspect -t 8 /path/to/kallisto/index.idx
```

Sample output:

```
[index] k-mer length: 31
[index] number of targets: 252,301
[index] number of k-mers: 155,644,518
[index] number of distinguishing flanking k-mers: 7,425,493
[inspect] Index version number = 12
[inspect] number of unitigs = 9411252
```

```
[inspect] minimizer length = 23
[inspect] max EC size = 3873
[inspect] number of ECs discarded = 0
```

##### 1.5 kallisto version

Prints out the version of the kallisto software that is being used

##### 1.6 kallisto cite

Prints out citation information

#### 2. bustools

bustools is run on BUS files generated by the `kallisto bus` command. The first step in working with BUS files is usually to sort the BUS file using `bustools sort`. This will organize the BUS file, making it suitable for use with other bustools commands. In a standard workflow, the sorted BUS file is error-corrected to a barcode on list via `bustools correct`, then sorted again, then quantified into count matrices via `bustools count`. There are many bustools commands, some of which are outside the scope of this protocol and some of which are in development, therefore only the bustools commands relevant to most RNA-seq analyses are presented here.

Many of the bustools commands can read from the standard input (stdin), by specifying `-` as the input file and write to standard output (stdout) using the `-p` flag if available.

##### 2.1 bustools sort

Sorts a BUS file. `bustools sort` (using the default options) should always be done before any additional processing of the BUS file following generation of the BUS file from the `kallisto bus` command. Many bustools commands will not work properly with an unsorted BUS file. Increasing the number of threads and maximum memory will speed up sorting.

The default behavior is to sort by barcode, UMI, equivalence class (ec), then the flag column.

Usage: bustools sort [options] bus-files

```
Arguments:
-t, --threads=INT         Number of threads to use (default: 1)
```

```
-m, --memory=STRING       Maximum memory used (default: 4G)

-T, --temp=STRING         Location and prefix for temporary files
                          (required if using -p, otherwise
                          defaults to output)

-o, --output=STRING       Filename to output sorted BUS file into

-p, --pipe                Write to standard output


    --umi                 Sort by UMI, barcode, then ec

    --count               Sort by multiplicity (count), barcode, UMI, then ec

    --flags               Sort by flag, ec, barcode, then UMI

    --flags-bc            Sort by flag, barcode, UMI, then ec

    --no-flags            Ignore and reset the flag column while sorting.
                          If read numbers are present in the flag column of
                          the BUS file, sorting using this option renders
                          BUS file suitable for use in generating
                          count matrices.
```

#### 2.2 bustools correct

Error-corrects the barcodes in a BUS file to an "on list".

Error correction is done based on a hamming distance 1 mismatch between each BUS file barcode sequence and each "on list" sequence. For barcode error correction, the "on list" file simply contains a list of sequences in the "on list".

Another operation supported is the replacement operation: Each "on list" sequence (in the first column of the "on list" file) has a replacement sequence (in the second column of the "on list" file) designated therefore if a BUS file barcode has an exact match to one of those "on list" sequences, it is replaced with its replacement sequence.

Note: The input BUS file need not be sorted.

Usage: bustools correct [options] bus-files

```
Arguments:
-o, --output=STRING      Filename to output barcode-corrected BUS file into

-w, --onlist=FILE        File containing the “on list” sequences

-p, --pipe               Write to standard output

-r, --replace            Perform the replacement operation rather than the
                         barcode error correction operation for the file
                         supplied in the -w option
```

#### 2.3 bustools count

Generates count matrices from BUS files that have been sorted and barcode-error-corrected.

Usage: bustools count [options] sorted-bus-files

```
Arguments:
-o, --output=STRING      The prefix of the output files for count matrices

-g, --genemap=FILE       File for mapping transcripts to genes
                         (when using kb ref in kb-python, this is the
                         t2g.txt file produced by kb ref)

-e, --ecmap=FILE         File for mapping equivalence classes to transcripts

-t, --txnames=FILE       File with names of transcripts

    --genecounts         Aggregate counts to genes only.
                         This option generates a gene count matrix; if this
                         option is not supplied, a transcript-compatibility
                         counts (TCC) matrix (where each equivalence class
                         gets a count) is generated instead.

    --umi-gene           Handles cases of UMI collisions. For example, a case
                         may be where two reads with the same UMI
                         sequence and the same barcode map to different
                         genes. With this option enabled, those reads are
                         considered to be two distinct molecules which were
                         unintentionally labeled with the same UMI, and hence
                         each gene gets a count.
```

| | |
|---|---|
| `--cm` | Counts multiplicities rather than UMIs. In other words, no UMI collapsing is performed and each mapped read is its own unique molecule regardless of the UMI sequence (i.e. the UMI sequence is ignored). |
| `-m, --multimapping` | Include bus records that map to multiple genes. When --genecounts is enabled, this option causes counts to be distributed uniformly across all the mapped genes (for example, if a read multimaps to two genes, each gene will get a count of 0.5). |
| `-s, --split=FILE` | Split output matrix in two (plus ambiguous) based on the list of transcript names supplied in this file. If a UMI (after collapsing) or a read maps to transcripts found in this file, the count is entered into a matrix file with the extension .2.mtx; if it maps to transcripts not in this file, the count is entered into a separate matrix file with the extension .mtx; if it maps to some transcripts in this file and some transcripts not in this file, the count is entered into a third matrix file with the extension .ambiguous.mtx.<br>When quantifying nascent, ambiguous, and mature RNA species, the nascent transcript names (which will actually simply be the gene IDs themselves) will be listed in the file supplied to --split so that the .mtx file contains the mature RNA counts, the .2.mtx file contains the nascent RNA counts, and the .ambiguous.mtx file contains the ambiguous RNA counts. Note that kb-python renames .mtx to .mature.mtx and renames 2.mtx to .nascent.mtx. |

Output:

Each output file is prefixed with what is supplied to the `--output` option. In kb count within kb-python, the prefix is cells_x_genes. Thus, the files outputted (when generating a gene count matrix via --genecounts) will be cells_x_genes.mtx (the matrix file), cells_x_genes.barcodes.txt (the barcodes; i.e. the rows of the matrix), and cells_x_genes.genes.txt (the genes; i.e. the columns of the matrix). When generating a TCC matrix, cells_x_genes.ec.txt will be generated in lieu of cells_x_genes.genes.txt as the columns of the matrix will be equivalence classes (ECs) rather than genes. If both sample-specific barcodes and cell barcodes are supplied (as is the case when one uses `--batch-barcodes` in `kallisto bus`), then an additional cells_x_genes.barcodes.prefix.txt file will be created containing the sample-specific barcodes.

The lines of this file correspond to the lines in the cells_x_genes.barcodes.txt (both files will have the same number of lines). Finally, when `--split` is supplied, additional .mtx matrix files will be generated (see the `--split` option described above).

#### 2.4 bustools inspect

Produces a report summarizing the contents of a sorted BUS file. The report can be output either to standard output or to a JSON file.

Usage: bustools inspect [options] sorted-bus-file

```
Arguments:
-o, --output=STRING       Filename to output sorted BUS file into

-e, --ecmap=FILE          File for mapping equivalence classes to transcripts

-w, --onlist=FILE         File containing the barcodes "on list"

-p, --pipe                Write to standard output
```

Sample report output in standard output (using `-p`):

```
Read in 3148815 BUS records
Total number of reads: 3431849

Number of distinct barcodes: 162360
Median number of reads per barcode: 1.000000
Mean number of reads per barcode: 21.137281

Number of distinct UMIs: 966593
Number of distinct barcode-UMI pairs: 3062719
Median number of UMIs per barcode: 1.000000
Mean number of UMIs per barcode: 18.863753

Estimated number of new records at 2x sequencing depth: 2719327

Number of distinct targets detected: 70492
Median number of targets per set: 2.000000
Mean number of targets per set: 3.091267

Number of reads with singleton target: 1233940

Estimated number of new targets at 2x seuqencing depth: 6168
```

```
Number of barcodes in agreement with on-list: 92889 (57.211752%)
Number of reads with barcode in agreement with on-list: 3281671
(95.623992%)
```

Sample report output in JSON format:


```
{
	"numRecords": 3148815,
	"numReads": 3431849,
	"numBarcodes": 162360,
	"medianReadsPerBarcode": 1.000000,
	"meanReadsPerBarcode": 21.137281,
	"numUMIs": 966593,
	"numBarcodeUMIs": 3062719,
	"medianUMIsPerBarcode": 1.000000,
	"meanUMIsPerBarcode": 18.863753,
	"gtRecords": 2719327,
	"numTargets": 70492,
	"medianTargetsPerSet": 2.000000,
	"meanTargetsPerSet": 3.091267,
	"numSingleton": 1233940,
	"gtTargets": 6168,
	"numBarcodesOnOnlist": 92889,
	"percentageBarcodesOnOnlist": 0.57211752,
	"numReadsOnOnlist": 3281671,
	"percentageReadsOnOnlist": 0.95623992
}
```


Note: The numTargets, medianTargetsPerSet, meanTargetsPerSet, numSingleton, and gtTargets values are only generated if the `--ecmap` option is provided. The numBarcodesOnOnlist, percentageBarcodesOnOnlist, numReadsOnOnlist, percentageReadsOnOnlist values are only generated if the `--onlist` is provided.

#### 2.5 bustools allowlist

Generates an “on list” based on the barcodes in a sorted BUS file. This is a way of generating an on list that the barcodes in the BUS file will be corrected to, for technologies that don’t provide an on list.

Usage: bustools allowlist [options] sorted-bus-file

```
Arguments:
-o, --output=STRING          Filename to output the "on list" into

-f, --threshold=INT          A highly optional parameter specifying the
                             minimum number of times a barcode must appear
                             to be included in "on list". If not provided, a
                             threshold will be determined based on the first
                             200 to 100200 BUS records.
```

#### 2.6 bustools capture

Separates a BUS file into multiple files according to the capture criteria.

Usage: bustools capture [options] bus-files

```
Capture options:
-F, --flags               Capture list is a list of flags to capture

-s, --transcripts         Capture list is a list of transcripts to capture

-u, --umis                Capture list is a list of UMI sequences to capture

-b, --barcode             Capture list is a list of barcodes to capture

Arguments:
-o, --output=STRING       Name of file for the captured BUS output

-x, --complement          Take complement of captured set
                          (i.e. output all BUS records that do NOT
                          match an entry in the capture list)

-c, --capture=FILE        File containing the "capture list"
                          (i.e. list of transcripts, transcripts, flags,
                          UMI sequences, or barcode sequences)

-e, --ecmap=FILE          File for mapping equivalence classes to
                          transcripts (required for --transcripts)

-t, --txnames=FILE        File with names of transcripts
                          (required for --transcripts)

-p, --pipe                Write to standard output
```

Note: If you use the `-b` (`--barcode`) option and want to capture all records containing a sample-specific barcode from running `--batch-barcodes` in `kallisto bus`, in the "capture list" file, enter the 16-bp sample-specific barcode followed by a * character (e.g. AAAAAAAAAAAAAACT*).

#### 2.7 bustools text

Converts a binary BUS file into its plaintext representation. The plaintext will have the columns (in order): barcode, UMI, equivalence class, count, flag, and pad. (Note: The last two columns will only be outputted if the respective option is specified by the user).

Usage: bustools text [options] bus-files

```
Arguments:
-o, --output=STRING       Filename of the output text file

-f, --flags               Write the flag column

-d, --pad                 Write the pad column
                          (the "pad" column is an additional 32-bit field
                          in the BUS file, in case one would like to use the
                          BUS format to store additional data for each BUS
                          record; this column is typically not used.)

-p, --pipe                Write to standard output

-a, --showAll             Show all 32 bases in the barcodes field (e.g. if
                          --batch-barcodes is specified in kallisto bus, the
                          cell barcodes are stored in barcodes field and are
                          used for bustools barcode correction to an on-list;
                          however, the artificial sample-specific barcodes
                          are stored as an additional "hidden" field in the
                          barcodes column, immediately preceding the cell
                          barcodes, and may be truncated or left-padded with
                          A's to fill the 32 bases. For example, if the cell
                          barcode is 12 bases, there will be 4 A's followed
                          by the 16-bp sample-specific barcode followed by
                          the 12-base cell barcode. If the cell barcode is 26
                          bases, the last 6 bases of the sample-specific
                          barcode will be shown followed by the 26-base cell
                          barcode).
```

An example of the plaintext output of a BUS file (with the flag column):

```
AAAAGATCACTATGCACTATCATC GCAAAACCTT     156  2    0
AAAAGATCAGATCGCACACTTTCA TAGAGTAACC     438  3    0
AAAAGATCAGATCGCAGCTCTACT TTAGGTATAG     1808 1    0
AAAAGATCAGCACCTCCTGACTTC AATCGGCATT     4481 1    0
```

If one runs `kallisto bus` with the `-n` (`--num`) option, the read number (zero-indexed) of the mapped reads will be stored in the flags column (i.e. the fifth column). One can view those read numbers using `bustools text` to identify which reads in the input FASTQ files mapped (and which reads were unmapped).

#### 2.8 bustools fromtext

Converts a plaintext representation of a BUS file to a binary BUS file. The plaintext input file should have four columns: barcode, UMI, equivalence class, and count. Optionally, a fifth column (the flags column) can be supplied.

Usage: bustools fromtext [options] text-files

```
Arguments:
-o, --output=STRING          Filename to write the output BUS file

-p, --pipe                   Write to standard output
```

#### 2.9 bustools extract

Extracts the successfully mapped sequencing reads from the input FASTQ files that were processed with `kallisto bus` with the `-n` (`--num`) option, which places the read number (zero-indexed) in the flags column of the BUS file. Although BUS files with read numbers present in the flags column should not be used for downstream quantification, they can be used by `bustools extract` to extract the original sequencing reads (as well as by `bustools text` to view the sequencing read number along with the barcode, UMI, and equivalence class).

Note: The BUS file must be sorted by flag. The output BUS file directly from kallisto should already be sorted by flag, but, if not, one can use apply `bustools sort --flag` on the BUS file.

Usage: bustools extract [options] sorted-by-flag-bus-file

```
Arguments:
-o, --output=STRING   Directory that the output FASTQ files will be stored in

-f, --fastq=STRING    FASTQ file(s) from which to extract reads
                      (comma-separated list). These should be the same files
                      used as input to kallisto bus.

-N, --nFastqs=INT     Number of FASTQ file(s) per run. For example, in 10xv3
                      where there are two FASTQ files (and R1 and R2 file),
                      --nFastqs=2 should be set.
```

This is especially useful to use in conjunction with `bustools capture` when one wishes to extract specific reads (e.g. reads that contain a certain barcode or reads whose equivalence class contains a certain transcript). Below, we show an example of how to extract reads from two input files: R1.fastq.gz and R2.fastq.gz entered into a kallisto bus run with results outputted into a directory named output_dir. We'll extract reads that are compatible with either the transcript ENSMUST00000171143.2 or ENSMUST00000131532.2.

Create a file called capture.txt containing the following two lines:

```
ENSMUST00000171143.2
ENSMUST00000131532.2
```

Run the following:

```
bustools capture -c capture.txt --transcripts \
--ecmap=output_dir/matrix.ec \
--txnames=output_dir/transcripts.txt -p \
output_dir/output.bus | bustools extract --nFastqs=2 \
--fastq=R1.fastq.gz,R2.fastq.gz -o extracted_output -
```

The capture results are directly piped into the extract command, and the extracted FASTQ sequencing reads output are placed into the paths extracted_output/1.fastq.gz and extracted_output/2.fastq.gz (for the input files R1.fastq.gz and R2.fastq.gz, respectively).

`bustools extract` does not work when you have sample-specific barcodes in your BUS file because each sample's read number (as recorded in the flags column of the BUS file) starts from 0. To work around this, you should first use `bustools capture` to isolate a specific sample and then supply that specific sample's FASTQ file(s).

#### 2.10 bustools umicorrect

Implements the UMI correction algorithm of UMI-tools and outputs a BUS file with the corrected UMIs.

Usage: bustools umicorrect [options] sorted-bus-file

```
Arguments:
-o, --output=STRING      Filename of the output BUS file with UMIs corrected

-p, --pipe               Write to standard output

-g, --genemap=FILE       File for mapping transcripts to genes
                         (when using kb ref in kb-python, this is the
                         t2g.txt file produced by kb ref)

-e, --ecmap=FILE         File for mapping equivalence classes to transcripts

-t, --txnames=FILE       File with names of transcripts
```

#### 2.11 bustools compress

Takes in a BUS file, sorted by barcode-umi-ec (i.e. the default option for `bustools sort`), and compresses it.

Usage: bustools compress [options] sorted-bus-file

```
Arguments:
-N, --chunk-size=INT           Number of rows to compress as a single block

-o, --output=STRING            Filename for the output compressed BUS file

-p, --pipe                     Write to standard output
```

#### 2.12 bustools decompress

Takes in a compressed BUS file and inflates (i.e. decompresses) it.

Usage: bustools decompress [options] compressed-bus-file

```
Arguments:
-o, --output=STRING            Filename for the output decompressed BUS file

-p, --pipe                     Write to standard output
```

#### 2.13 bustools version

Prints out the version of the bustools software that is being used.

#### 2.14 bustools cite

Prints out citation information.

**Supplementary Tutorial**: An example mouse multiplexed single-nucleus SPLiT-seq preprocessing workflow.

Here we describe how to process a mouse multiplexed single-nucleus SPLiT-seq assay. The input FASTQ files are split across multiple subpools such that two cells may have the same cell barcode but be in different subpools. The SPLiT-seq assay uses both oligo-dT and random hexamer primers (which are represented in the third component of the cell barcode, corresponding to the first round of split pooling). As a result, two sets of matrices will be produced: One with both the oligo-dT and random hexamer barcodes in the same count matrix and one with the oligo-dT barcodes converted into the random hexamer barcodes (so that each barcode is unique to one nucleus). This facilitates investigation of each library type separately (should one wish to generate an “oligo-dT” count matrix and a “random hexamer” count matrix) as well as of the two library types combined together.

**1. Install kb-python.**

```
pip install kb_python
```

**2. Download the mouse genome and annotation files.**

```
wget ftp.ensembl.org/pub/release-108/fasta/mus_musculus/dna/Mus_musculus.GRCm39.dna.primary_assembly.fa.gz

wget ftp.ensembl.org/pub/release-108/gtf/mus_musculus/Mus_musculus.GRCm39.108.gtf.gz
```

**3. Build the index.**

To illustrate index generation with GTF filtering we show below how to filter the GTF file to only keep the relevant biotypes (the same ones that are used in the CellRanger reference). This can improve both accuracy and efficiency. Additional methods to optimize the GTF file can also be used such as the one proposed in Pool et al., 2023[39] which can greatly increase gene detection sensitivity.

```
kb ref --workflow=nac -i index.idx -g t2g.txt \
    -c1 cdna.txt -c2 nascent.txt -f1 cdna.fasta -f2 nascent.fasta \
    --include-attribute gene_biotype:protein_coding \
    --include-attribute gene_biotype:lncRNA \
    --include-attribute gene_biotype:lincRNA \
    --include-attribute gene_biotype:antisense \
    --include-attribute gene_biotype:IG_LV_gene \
    --include-attribute gene_biotype:IG_V_gene \
    --include-attribute gene_biotype:IG_V_pseudogene \
    --include-attribute gene_biotype:IG_D_gene \
    --include-attribute gene_biotype:IG_J_gene \
    --include-attribute gene_biotype:IG_J_pseudogene \
    --include-attribute gene_biotype:IG_C_gene \
    --include-attribute gene_biotype:IG_C_pseudogene \
    --include-attribute gene_biotype:TR_V_gene \
    --include-attribute gene_biotype:TR_V_pseudogene \
    --include-attribute gene_biotype:TR_D_gene \
    --include-attribute gene_biotype:TR_J_gene \
    --include-attribute gene_biotype:TR_J_pseudogene \
    --include-attribute gene_biotype:TR_C_gene \
    Mus_musculus.GRCm39.dna.primary_assembly.fa.gz \
    Mus_musculus.GRCm39.108.gtf.gz
```

**4. Map the input sequencing reads to the index.**

This assay has multiple FASTQ files across multiple subpools as well as two primer types. To process this, we supply a batch.txt file containing the FASTQ files along with their designated subpool, a barcodes.txt file containing the three barcode components (since the assay contains three 8-bp barcodes, each separated by a linker, in the first read file), and a replace.txt file designating how to convert the random hexamer barcodes to the oligo-dT barcodes for the "combined" matrix. The final command to run with these files is as follows:

```
kb count --strand=forward -r replace.txt -w barcodes.txt \
     --workflow=nac -i index.idx -g t2g.txt -c1 cdna.txt \
     -c2 nascent.txt -x 1,10,18,1,48,56,1,78,86:1,0,10:0,0,0 \
     --sum=total -o output_dir --batch-barcodes batch.txt
```

**5. Analyze the output.**

Output (both the oligo-dT and random hexamer barcodes in the same count matrix):

- output_dir/counts_unfiltered/cells_x_genes.mature.mtx
- output_dir/counts_unfiltered/cells_x_genes.nascent.mtx
- output_dir/counts_unfiltered/cells_x_genes.ambiguous.mtx
- output_dir/counts_unfiltered/cells_x_genes.cell.mtx
- output_dir/counts_unfiltered/cells_x_genes.nucleus.mtx
- output_dir/counts_unfiltered/cells_x_genes.total.mtx
- output_dir/counts_unfiltered/cells_x_genes.barcodes.txt
- output_dir/counts_unfiltered/cells_x_genes.barcodes.prefix.txt
- output_dir/counts_unfiltered/cells_x_genes.genes.txt
- output_dir/counts_unfiltered/cells_x_genes.genes.names.txt

Output (the oligo-dT and random hexamer barcodes are combined):

- output_dir/counts_unfiltered_modified/cells_x_genes.mature.mtx
- output_dir/counts_unfiltered_modified/cells_x_genes.nascent.mtx
- output_dir/counts_unfiltered_modified/cells_x_genes.ambiguous.mtx
- output_dir/counts_unfiltered_modified/cells_x_genes.cell.mtx
- output_dir/counts_unfiltered_modified/cells_x_genes.nucleus.mtx
- output_dir/counts_unfiltered_modified/cells_x_genes.total.mtx
- output_dir/counts_unfiltered_modified/cells_x_genes.barcodes.txt
- output_dir/counts_unfiltered_modified/cells_x_genes.barcodes.prefix.txt
- output_dir/counts_unfiltered_modified/cells_x_genes.genes.txt
- output_dir/counts_unfiltered_modified/cells_x_genes.genes.names.txt

Note that the cells_x_genes.barcodes.prefix.txt will contain a unique identifier for each subpool.

**Information about batch.txt, barcodes.txt, and replace.txt files:**

**batch.txt:**

Example with three subpools, each sequenced on four lanes:

```
subpool_1    S1_lane1_R1.fastq.gz    S1_lane1_R2.fastq.gz
subpool_1    S1_lane2_R1.fastq.gz    S1_lane2_R2.fastq.gz
subpool_1    S1_lane3_R1.fastq.gz    S1_lane3_R2.fastq.gz
subpool_1    S1_lane4_R1.fastq.gz    S1_lane4_R2.fastq.gz
subpool_2    S2_lane1_R1.fastq.gz    S2_lane1_R2.fastq.gz
subpool_2    S2_lane2_R1.fastq.gz    S2_lane2_R2.fastq.gz
subpool_2    S2_lane3_R1.fastq.gz    S2_lane3_R2.fastq.gz
subpool_2    S2_lane4_R1.fastq.gz    S2_lane4_R2.fastq.gz
subpool_3    S3_lane1_R1.fastq.gz    S3_lane1_R2.fastq.gz
subpool_3    S3_lane2_R1.fastq.gz    S3_lane2_R2.fastq.gz
subpool_3    S3_lane3_R1.fastq.gz    S3_lane3_R2.fastq.gz
subpool_3    S3_lane4_R1.fastq.gz    S3_lane4_R2.fastq.gz
```

In this configuration, subpool_1 will have the sample-specific barcode AAAAAAAAAAAAAAAA, subpool_2 will have the sample-specific barcode AAAAAAAAAAAAAAAC, and subpool_3 will have the sample-specific barcode AAAAAAAAAAAAAAAG. This mapping can be found in the output_dir/matrix.cells and output_dir/matrix.sample.barcodes files. These sample-specific barcodes are found in cells_x_genes.barcodes.prefix.txt to identify the subpool a specific cell barcode originated from when inspecting the count matrices.

**barcodes.txt:**

The cell barcodes contain three 8-bp components so we should correct each component individually to its own “on list”. This can be done by having multiple columns in the barcodes.txt file. Note that the first two columns have 96 barcodes and the third column has 192 barcodes.

```
AACGTGAT AACGTGAT CATTCCTA
AAACATCG AAACATCG CTTCATCA
ATGCCTAA ATGCCTAA CCTATATC
AGTGGTCA AGTGGTCA ACATTTAC
ACCACTGT ACCACTGT ACTTAGCT
ACATTGGC ACATTGGC CCAATTCT
CAGATCTG CAGATCTG GCCTATCT
CATCAAGT CATCAAGT ATGCTGCT
CGCTGATC CGCTGATC CATTTACA
ACAAGCTA ACAAGCTA ACTCGTAA
CTGTAGCC CTGTAGCC CCTTTGCA
```

| | | |
|---|---|---|
| AGTACAAG | AGTACAAG | ACTCCTGC |
| AACAACCA | AACAACCA | ATTTGGCA |
| AACCGAGA | AACCGAGA | TTATTCTG |
| AACGCTTA | AACGCTTA | TCATGCTC |
| AAGACGGA | AAGACGGA | CATACTTC |
| AAGGTACA | AAGGTACA | CCGTTCTA |
| ACACAGAA | ACACAGAA | GCTTCATA |
| ACAGCAGA | ACAGCAGA | CTCTGTGC |
| ACCTCCAA | ACCTCCAA | CCCTTATA |
| ACGCTCGA | ACGCTCGA | ACTGCTCT |
| ACGTATCA | ACGTATCA | CTCTAATC |
| ACTATGCA | ACTATGCA | ACCCTTGC |
| AGAGTCAA | AGAGTCAA | ATCTTAGG |
| AGATCGCA | AGATCGCA | CATGTCTC |
| AGCAGGAA | AGCAGGAA | TCATTGCA |
| AGTCACTA | AGTCACTA | ACACCTTT |
| ATCCTGTA | ATCCTGTA | AATTTCTC |
| ATTGAGGA | ATTGAGGA | ATTCATGG |
| CAACCACA | CAACCACA | ACTTTACC |
| GACTAGTA | GACTAGTA | CTTCTAAC |
| CAATGGAA | CAATGGAA | CTATTTCA |
| CACTTCGA | CACTTCGA | TCTCATGC |
| CAGCGTTA | CAGCGTTA | ATCCTTAC |
| CATACCAA | CATACCAA | TAAATATC |
| CCAGTTCA | CCAGTTCA | TTACCTGC |
| CCGAAGTA | CCGAAGTA | CACTTTCA |
| CCGTGAGA | CCGTGAGA | CACCTTTA |
| CCTCCTGA | CCTCCTGA | CTGACTTC |
| CGAACTTA | CGAACTTA | CATTTGGA |
| CGACTGGA | CGACTGGA | GCTCTACT |
| CGCATACA | CGCATACA | GTTACGTA |
| CTCAATGA | CTCAATGA | CCTGTTGC |
| CTGAGCCA | CTGAGCCA | CTATCATC |
| CTGGCATA | CTGGCATA | GCTATCAT |
| GAATCTGA | GAATCTGA | ACATTCAT |
| CAAGACTA | CAAGACTA | TTCGCTAC |
| GAGCTGAA | GAGCTGAA | CATTCTAC |
| GATAGACA | GATAGACA | CACTTATC |
| GCCACATA | GCCACATA | ATAAGCTC |
| GCGAGTAA | GCGAGTAA | TCATCCTG |
| GCTAACGA | GCTAACGA | CCTGGTAT |
| GCTCGGTA | GCTCGGTA | TGGTATAC |
| GGAGAACA | GGAGAACA | TTGGGAGA |
| GGTGCGAA | GGTGCGAA | ACTTCATC |

| | | |
|---|---|---|
| GTACGCAA | GTACGCAA | TCTCTAGC |
| GTCGTAGA | GTCGTAGA | ATGCCCTT |
| GTCTGTCA | GTCTGTCA | CCCAATTT |
| GTGTTCTA | GTGTTCTA | ACTATATA |
| TAGGATGA | TAGGATGA | CTCTATAC |
| TATCAGCA | TATCAGCA | CTGTCTCA |
| TCCGTCTA | TCCGTCTA | GACCTTTC |
| TCTTCACA | TCTTCACA | GATTTGGC |
| TGAAGAGA | TGAAGAGA | CGTCTAGG |
| TGGAACAA | TGGAACAA | TACTCGAA |
| TGGCTTCA | TGGCTTCA | CAGCCTTT |
| TGGTGGTA | TGGTGGTA | CCTCATTA |
| TTCACGCA | TTCACGCA | CTTATACC |
| AACTCACC | AACTCACC | TCTATTAC |
| AAGAGATC | AAGAGATC | CCTGCATT |
| AAGGACAC | AAGGACAC | CAATCCTT |
| AATCCGTC | AATCCGTC | TTGTCTTA |
| AATGTTGC | AATGTTGC | TCACTTTA |
| ACACGACC | ACACGACC | TGCTTGGG |
| ACAGATTC | ACAGATTC | CGCTCATT |
| AGATGTAC | AGATGTAC | GCCTCTAT |
| AGCACCTC | AGCACCTC | GAGCACAA |
| AGCCATGC | AGCCATGC | CTCTTAAC |
| AGGCTAAC | AGGCTAAC | TCTAGGCT |
| ATAGCGAC | ATAGCGAC | AATTCTGC |
| ATCATTCC | ATCATTCC | CATTCTCA |
| ATTGGCTC | ATTGGCTC | ACTTGCCT |
| CAAGGAGC | CAAGGAGC | ATCATTGC |
| CACCTTAC | CACCTTAC | GTTCAACA |
| CCATCCTC | CCATCCTC | CCATTTGC |
| CCGACAAC | CCGACAAC | GACTTTGC |
| CCTAATCC | CCTAATCC | ATTGGCTC |
| CCTCTATC | CCTCTATC | GTGCTAGC |
| CGACACAC | CGACACAC | CTTTCAAC |
| CGGATTGC | CGGATTGC | ACTATTGC |
| CTAAGGTC | CTAAGGTC | ACTGGCTT |
| GAACAGGC | GAACAGGC | ATTAGGCT |
| GACAGTGC | GACAGTGC | GCCTTTCA |
| GAGTTAGC | GAGTTAGC | ATTCTAGG |
| GATGAATC | GATGAATC | CCTTACAT |
| GCCAAGAC | GCCAAGAC | ACATTTGG |
| - | - | CATCATCC |
| - | - | CTGCTTTG |
| - | - | CTAAGGGA |

| | | |
|---|---|---|
| - | - | GCTTATAG |
| - | - | TCTGATCC |
| - | - | TCTCTTGG |
| - | - | CAATTTCC |
| - | - | AGTCTCTT |
| - | - | TGCTGCTC |
| - | - | GTATTTCC |
| - | - | TTCCTGTG |
| - | - | GCTGCTTC |
| - | - | TATGTGTC |
| - | - | CAATTCTC |
| - | - | TGGTCTCC |
| - | - | GCTCTTTA |
| - | - | GCTGCATG |
| - | - | ACTCATTT |
| - | - | AGTCTTGG |
| - | - | GGTTCTTC |
| - | - | TCATGTTG |
| - | - | ATTTTGCC |
| - | - | CTTCTGTA |
| - | - | GTCCATCT |
| - | - | GCTATCTC |
| - | - | TAGTTTCC |
| - | - | TCCATTAT |
| - | - | AGGATTAA |
| - | - | AATCTTTC |
| - | - | GTCATATG |
| - | - | GTGCTTCC |
| - | - | ATGTGTTG |
| - | - | CCATCTTG |
| - | - | TACTGTCT |
| - | - | TTCATCGC |
| - | - | ACTGTGGG |
| - | - | TCTGTGCC |
| - | - | TCAATCTC |
| - | - | GTCCTCTG |
| - | - | TTACATTC |
| - | - | ATTCTGTC |
| - | - | TGTGTATG |
| - | - | TCCATTTG |
| - | - | TTAGCTTC |
| - | - | GTGCTTGA |
| - | - | GTTTGTGA |
| - | - | GAAATTAG |

| | | |
|---|---|---|
| - | - | GCAAATTC |
| - | - | GAGGTTGA |
| - | - | CCTGTCTG |
| - | - | GTGGGTTC |
| - | - | TTTGCATC |
| - | - | AGGTAATA |
| - | - | GTGCCTTC |
| - | - | ATGTTTCC |
| - | - | CTTAATTC |
| - | - | TCTGGCTC |
| - | - | CATCATTT |
| - | - | GTTGTCTC |
| - | - | ATCTTCTG |
| - | - | TGTTTGCC |
| - | - | TTCTGTCA |
| - | - | ACGGACTC |
| - | - | TTTGGTCA |
| - | - | TATCCGGG |
| - | - | TGTCATTC |
| - | - | ATTCTCTG |
| - | - | TGGCTTCC |
| - | - | TTGTTGCC |
| - | - | GTCATCTC |
| - | - | TTGCTCAT |
| - | - | CTGTCTGC |
| - | - | TATATTCC |
| - | - | ATATTGGC |
| - | - | GTGTCCTC |
| - | - | ATCTTCAT |
| - | - | CGTGGTTG |
| - | - | TTGCATCC |
| - | - | TCTTAATC |
| - | - | TGCATTTC |
| - | - | GATGTTTC |
| - | - | ATCTTGTC |
| - | - | TCATATTC |
| - | - | TGGCCTCT |
| - | - | CGTTGTCT |
| - | - | TCTTGTCA |
| - | - | TATTCCTG |
| - | - | TCCATGTC |
| - | - | TTGTCATC |
| - | - | ATTTCCTG |
| - | - | GTGTCTCC |

```
-         -         GTGTGTGT
-         -         TATGCTTC
-         -         ATGGTGTT
-         -         GAATAATG
-         -         CCTCTGTG
```

**replace.txt:**

This file contains the instructions on how to produce the "modified" count matrix in output_dir/counts_unfiltered_modified/ – the output directory which contains the combined oligo-dT and random hexamer barcodes wherein the random hexamer barcodes (first column of the file) are converted to their oligo-dT counterparts (second column of the file). These barcodes, being the third component of the barcode, occur at the end of the final barcode string. The asterisk (*) at the beginning of the replacement string tells bustools to convert the nucleotides at the end of the barcode sequence. As an example, the barcode sequence AACAACCATGAAGAGACATCATCC will be converted into AACAACCATGAAGAGACATTCCTA in the final output in the output_dir/counts_unfiltered_modified/ directory.

```
CATCATCC *CATTCCTA
CTGCTTTG *CTTCATCA
CTAAGGGA *CCTATATC
GCTTATAG *ACATTTAC
TCTGATCC *ACTTAGCT
TCTCTTGG *CCAATTCT
CAATTTCC *GCCTATCT
AGTCTCTT *ATGCTGCT
TGCTGCTC *CATTTACA
GTATTTCC *ACTCGTAA
TTCCTGTG *CCTTTGCA
GCTGCTTC *ACTCCTGC
TATGTGTC *ATTTGGCA
CAATTCTC *TTATTCTG
TGGTCTCC *TCATGCTC
GCTCTTTA *CATACTTC
GCTGCATG *CCGTTCTA
ACTCATTT *GCTTCATA
AGTCTTGG *CTCTGTGC
GGTTCTTC *CCCTTATA
TCATGTTG *ACTGCTCT
ATTTTGCC *CTCTAATC
CTTCTGTA *ACCCTTGC
GTCCATCT *ATCTTAGG
```

GCTATCTC *CATGTCTC
TAGTTTCC *TCATTGCA
TCCATTAT *ACACCTTT
AGGATTAA *AATTTCTC
AATCTTTC *ATTCATGG
GTCATATG *ACTTTACC
GTGCTTCC *CTTCTAAC
ATGTGTTG *CTATTTCA
CCATCTTG *TCTCATGC
TACTGTCT *ATCCTTAC
TTCATCGC *TAAATATC
ACTGTGGG *TTACCTGC
TCTGTGCC *CACTTTCA
TCAATCTC *CACCTTTA
GTCCTCTG *CTGACTTC
TTACATTC *CATTTGGA
ATTCTGTC *GCTCTACT
TGTGTATG *GTTACGTA
TCCATTTG *CCTGTTGC
TTAGCTTC *CTATCATC
GTGCTTGA *GCTATCAT
GTTTGTGA *ACATTCAT
GAAATTAG *TTCGCTAC
GCAAATTC *CATTCTAC
GAGGTTGA *CACTTATC
CCTGTCTG *ATAAGCTC
GTGGGTTC *TCATCCTG
TTTGCATC *CCTGGTAT
AGGTAATA *TGGTATAC
GTGCCTTC *TTGGGAGA
ATGTTTCC *ACTTCATC
CTTAATTC *TCTCTAGC
TCTGGCTC *ATGCCCTT
CATCATTT *CCCAATTT
GTTGTCTC *ACTATATA
ATCTTCTG *CTCTATAC
TGTTTGCC *CTGTCTCA
TTCTGTCA *GACCTTTC
ACGGACTC *GATTTGGC
TTTGGTCA *CGTCTAGG
TATCCGGG *TACTCGAA
TGTCATTC *CAGCCTTT
ATTCTCTG *CCTCATTA
TGGCTTCC *CTTATACC

```
TTGTTGCC *TCTATTAC
GTCATCTC *CCTGCATT
TTGCTCAT *CAATCCTT
CTGTCTGC *TTGTCTTA
TATATTCC *TCACTTTA
ATATTGGC *TGCTTGGG
GTGTCCTC *CGCTCATT
ATCTTCAT *GCCTCTAT
CGTGGTTG *GAGCACAA
TTGCATCC *CTCTTAAC
TCTTAATC *TCTAGGCT
TGCATTTC *AATTCTGC
GATGTTTC *CATTCTCA
ATCTTGTC *ACTTGCCT
TCATATTC *ATCATTGC
TGGCCTCT *GTTCAACA
CGTTGTCT *CCATTTGC
TCTTGTCA *GACTTTGC
TATTCCTG *ATTGGCTC
TCCATGTC *GTGCTAGC
TTGTCATC *CTTTCAAC
ATTTCCTG *ACTATTGC
GTGTCTCC *ACTGGCTT
GTGTGTGT *ATTAGGCT
TATGCTTC *GCCTTTCA
ATGGTGTT *ATTCTAGG
GAATAATG *CCTTACAT
CCTCTGTG *ACATTTGG
```

**The commands run by kb count in this example:**

```
mkdir -p output_dir/tmp

mkdir -p output_dir

kallisto bus -i index.idx -o output_dir -x
1,10,18,1,48,56,1,78,86:1,0,10:0,0,0 -t 8 --fr-stranded
--batch-barcodes --batch batch.txt

bustools sort -o output_dir/tmp/output.s.bus -T output_dir/tmp -t 8
-m 4G output_dir/output.bus
```

```
bustools inspect -o output_dir/inspect.json -w barcodes.txt
output_dir/tmp/output.s.bus

bustools correct -o output_dir/tmp/output.s.c.bus -w barcodes.txt
output_dir/tmp/output.s.bus

bustools sort -o output_dir/output.unfiltered.bus -T output_dir/tmp
-t 8 -m 4G output_dir/tmp/output.s.c.bus

mkdir -p output_dir/counts_unfiltered

bustools count -o output_dir/counts_unfiltered/cells_x_genes -g
t2g.txt -e output_dir/matrix.ec -t output_dir/transcripts.txt -s
nascent.txt --genecounts --umi-gene
output_dir/output.unfiltered.bus

mv output_dir/counts_unfiltered/cells_x_genes.mtx
output_dir/counts_unfiltered/cells_x_genes.mature.mtx

mv output_dir/counts_unfiltered/cells_x_genes.2.mtx
output_dir/counts_unfiltered/cells_x_genes.nascent.mtx

bustools correct -o output_dir/tmp/output.unfiltered.c.bus -w
replace.txt output_dir/output.unfiltered.bus --replace
bustools sort -o output_dir/output_modified.unfiltered.bus -T
output_dir/tmp -t 8 -m 4G output_dir/tmp/output.unfiltered.c.bus

mkdir -p output_dir/counts_unfiltered_modified

bustools count -o
output_dir/counts_unfiltered_modified/cells_x_genes -g t2g.txt -e
output_dir/matrix.ec -t output_dir/transcripts.txt -s nascent.txt
--genecounts --umi-gene output_dir/output_modified.unfiltered.bus

mv output_dir/counts_unfiltered_modified/cells_x_genes.mtx
output_dir/counts_unfiltered_modified/cells_x_genes.mature.mtx

mv output_dir/counts_unfiltered_modified/cells_x_genes.2.mtx
output_dir/counts_unfiltered_modified/cells_x_genes.nascent.mtx

rm -rf output_dir/tmp
```
